# A novel polycistronic method tailored for engineering split GECIs

**DOI:** 10.1101/2023.07.16.549202

**Authors:** Shunit Olszakier, Wessal Hussein, Ronit Heinrich, Michael Andreyanov, Yara Otor, Jackie Schiller, Shai Kellner, Shai Berlin

**Affiliations:** Department of Neuroscience, Ruth and Bruce Rappaport Faculty of Medicine, Technion- Israel Institute of Technology, Israel

## Abstract

We assessed the feasibility of using stop-codons as means to obtain polycistronic expression in eukaryotic cells. We show robust bicistronic expression of different open reading frames (ORFs), when these are cloned in-sequence and simply separated by stop codons (in-or out-of-frame), in heterologous expression systems and primary neurons. We further find this method to support polycistronic expression of three stop-codon-separated ORFs *in vivo*, which guided us to develop a technicolor Genetically-Encoded Functional Rainbow Indicators (GEFRIs) for monitoring cellular morphology and neuronal firing, concomitantly. These findings guided us to develop a new technique we denote *SPLIT*—<u>S</u>top-codon mediated <u>P</u>o<u>l</u>ycistronic <u>I</u>nduction in He<u>T</u>erologous expression systems— for rapid and easy development of fragmented proteins by the sole use of stop codons. We validated the *SPLIT* method by generating several new split-GFP variants, then engineer a palette of functional split-GCaMP6s variants and, lastly, generate a split ca^2+^-probe localized at ER and mitochondria junctions, denoted split-MEGIC. With the use of the probe, we show presence and activity of mito-ER contact sites within individual dendritic spines. Split-MEGIC can thereby be imaged by two-photon excitation *in vivo* in mice brains and, by standard confocal microscope in transgenic zebrafish larvae. Together, we explore non-canonical translation mechanisms and show these to be highly pervasive in various cell types *in vitro* and *in vivo*. We harness translation re-initiation to express multiple ORFs, to engineer rainbow indicators and to swiftly produce functional split-proteins and probes.

## Introduction

Split-proteins are purposefully fragmented proteins, typically into two fragments, that can spontaneously assemble and reconstitute the functional unit when jointly expressed^1^. A prototypical example is split-GFP^2^, in which instance the fragments are non-fluorescent on their own unless re-assembled^2–5^. Intuitively, split-proteins are particularly tailored for monitoring protein-protein interactions^6^, for instance by methods such as Bimolecular Fluorescence Complementation (BiFC^3^), however the range of biological activities that can be monitored using split-proteins is much broader. Indeed, split-GFP has also been leveraged for tracing neuronal connectivity^7,8^, tracking viral fusion, to determine membrane protein topology, to study protein-RNA interactions, report on protease activity, or used as antibody-like peptides for tagging proteins (see ^9^). Despite their vast potential (and despite emerging in the late 1950s’^10^), split proteins are a rare commodity, exemplified by the very few extant examples, such as split-ribonuclease^10^, split-ubiquitin^11^, split-biotin-ligase^12^ and, only recently, split-genome editing tools^13–15^ (see reviews^6,16^). We suggest that this deficiency persists because these are extremely laborious to engineer^2,4,9^; entailing the need for combining multiple methods, namely molecular, structural and/or computational methods (e.g., scrutiny of protein structure and sequence, followed by sequential cloning, protein partitioning and high-throughput screenings, not to mention protein evolution methods for fragment stabilization, etc.^2,4,12–19^). Regardless the method employed, trial-and-error remains the most popular route for developing split-proteins and, resultantly, is mostly limited to highly specialized research groups^20,21^.

We coveted easing this process with the aim to rapidly develop split-genetically encoded calcium indicators (GECIs)^22–24^. GECIs are powerful fluorescent tools commonly employed to interrogate neuronal activity^22,25,26^. We hypothesized that splitting the probe may provide additional means for selective and gated-expression, as well as bestow onto the indicator additional capabilities, for instance the ability to also serve as dual BiFC-reporter and calcium-indicator. This would extend the capabilities of the reporter, rendering it multifunctional, much like photoactivatable^27,28^ and photoconvertible^29^ schemes have yielded *functional highlighting* GECI-variants^30^. Notably, there are no *bona fide* split-GECIs^31^.

One major hurdle in the developing split-proteins is the need to randomly divide the coding sequence (i.e., open reading frame, ORF) of the protein and encode these within independent expression vectors, repeatedly, until a viable splitting site is discovered (e.g., ^12^). We coveted constraining the cloning process to a single vector, to ease and hasten the process; leading us to focus on methods that enable production of two proteins from the same vector, namely polycistronic expression. Of note, polycistronic expression has not been previously employed as a means for engineering split-proteins^21^. Briefly, polycistronic expression can be obtained by inserting a suite of unique sequences, commonly internal ribosome entry sites (IRES)^32^, self-cleavable peptides (e.g., P2A)^33,34^ or promoters^35,36^, that yield translation of two independent (i..e, separate) proteins. However, these are not suitable for our aim as these entail insertion of large sequences between the fragments, or yield protein products with superfluous residues, or products that do not undergo full separation, to name a few^32,37,38^. We consequently focused our attention on non-canonical translation mechanisms in mammalian cells that can produce two ORFs^1^ and, importantly, that require minimal perturbation (or additions) to the DNA. These narrowed our search onto mechanisms such as alternative initiation sites, re-initiation (REI), and leaky ribosome scanning (LRS)^39–41^. Briefly, REI is observed following short translation events (typically <100 base pairs; bp), during which the 40S subunit may encounter a secondary downstream AUG that promotes re-initiation^42–47^. Secondary translation events frequently transpire shortly after premature stop codons^44,48–50^; rarely after long ORFs^41,51–53^. LRS, on the other hand, entails a scenario whereby the ribosome travels past the primary start codon, without initiating translation, until it encounters an alternative start codon downstream of the primary methionine^54^. Notably, this process is less restricted by features such as distances from initial AUG (as in the case of REI), but is highly dependent on the context, namely on sequences in the vicinity of the primary and secondary start codons such as *Kozak* sequences^44^. A weak *kozak* sequence next to the primary AUG will potentially increase likelihood of LRS^40,46^. However, as in REI, LRS is mostly observed following short distances past the primary initiation site^44,55^.

To be applicable for splitting proteins, it remains to be determined whether non-canonical translation past stop codons can occur at larger distances past the primary AUG. Secondly, whether different cell types, notably neurons, can accommodate these mechanisms. Third, whether stop-codon mediated polycistronic expression is limited to two ORFs. Here we have directly examined the feasibility of using stop-codons as means for polycistronic expression and demonstrate that various cells, including neurons *in vitro* and *in vivo*, can readily translate up to three independent ORFs, cloned in-sequence and solely separated by stop codons. We then employ the technique to engineer new and seamless split genetically encoded probes for use *in vitro* and *in vivo*.

## Results

### Polycistronic expression by use of stop codons

To explore the feasibility, and propensity, of translation of two ORFs—without added promoters or sequences—by mammalian cells, we cloned a membrane-tethered SNAP^1^ (SNAP_mem_^58^) consisting of an extracellular-facing SNAP-tag^56^ along an intracellular red fluorescent protein (tdTomato; tdTom^59^); estimated at 75 kDa (**Fig. 1a** and **methods**). This clone displayed expression at internal membranes of human embryonic kidney cells (HEK293t, HEK in brief), mostly ER (**Suppl. 1a vs. Suppl. 1b**), as well as at plasma membrane assessed by impermeable surface-labeling of SNAP_mem_ (**Fig. 1a, surface labeling-green, magenta-protein expression, right panel**). We then introduced a single stop codon (UGA) immediately following the transmembrane domain, preceding tdTom (SNAP*tdTom; **Fig. 1b, cartoon**); placing the stop codon (and ensuing methionine of tdTom) 800 base-pairs (bps) following the primary methionine (i.e., initiation site). In HEK, we observed robust expression of tdTom (denoted ORF2, **Fig. 1b, micrograph**), along improvement in surface expression of the first ORF (ORF1- SNAP_mem_) (**Fig. 1b-d**). ORF1 and ORF2 were jointly detected in the majority (∼90%) of cells (**Fig. 1e**), and this phenomenon was similarly observed in HeLa cells (**Fig. 1e, bottom; Suppl. 2a**) and cultured primary hippocampal neurons (**Suppl. 2b**). Interestingly, fluorescence of ORF2 homogenously filled the cell (cytoplasm and nucleus), suggesting a soluble protein, as opposed to the membranes-restricted red fluorescence obtained by the parent clone (**Fig. 1b-c, and Suppl. 2a**), which we indeed confirmed by Western blot (**Fig. 1f**). Thus, these observations rule-out stop codon readthrough (RT)^60^ and show that polycistronic expression of two ORFs, solely separated by a single stop codon, is feasible and robust in multiple cell types. Moreover, these results deviate from previous reports by showing that expression of ORF2 can occur past greater distances (past 800 bps) than previously reported (by premature stop-codons, ∼100 bps^43,49^) (see **methods and sequences**) (and see ^61^).

**Figure 1.**
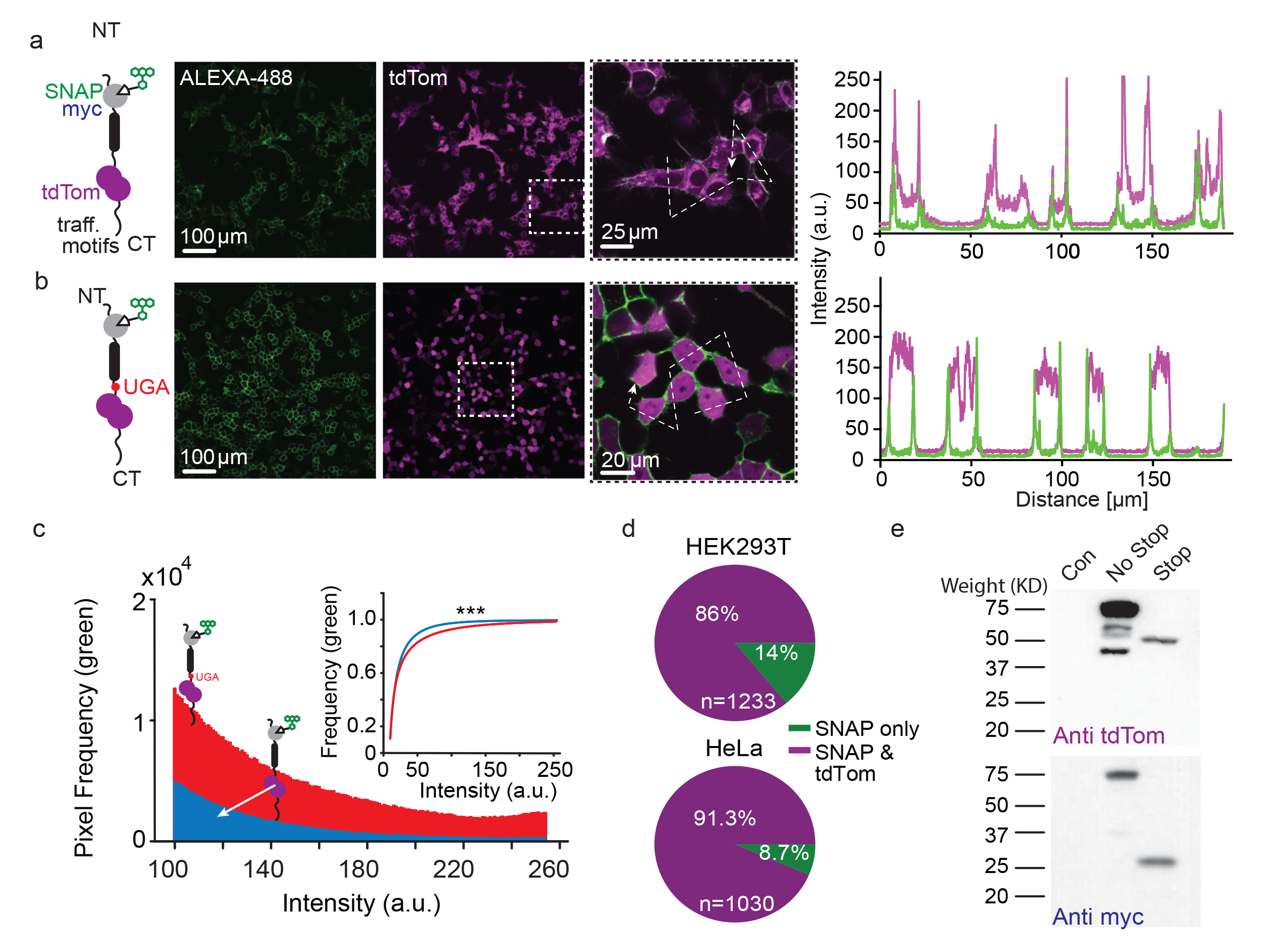
Bicistronic expression of two distinct ORFS obtained by a single stop codon. **a, b.** Illustration of SNAP-tdTom **(a)** and SNAP*tdTom **(b)** clones (left panels). The clones contain a SNAP-tag domain (grey) to conjugate a fluorescently-labeled substrate (benzylguanine, BG; white triangle and Alexa488, green), a myc-tag (blue), a PDGFR transmembrane domain (black), and a fluorescent reporter tdTom (magenta). The UGA stop codon is denoted by a red circle (in **b**). (middle panels) Micrographs of HEK293 cells expressing the SNAP-tdTom (**a**) and SNAP*tdTom (**b**) constructs (magenta channel) and labeled by BG-Alexa488 (green). Inset (dashed white region) depicts higher power micrographs showing diverging expression patterns of green and magenta, summarized in fluorescence profile plots (rightmost panels). **c.** Comparison of expression of the clones; assessed by measuring surface expression of BG-Alexa488. Plot shows pixel-pixel comparison of green BG-Alexa488 conjugated SNAP-tag fluorescence intensities of SNAP-tdTom construct (blue) and SNAP*tdTom (red),and normalized cumulative plot (inset). Kolmogorov-Smirnov comparison (***, p< 0.001) shows that the presence of stop codon yields significantly improved expression of SNAP-tag. **d.** Occurrence of cells showing expression of both ORFs (SNAP and tdTom, magenta area) vs. ORF1 (SNAP only, green area) in transfected HEK293t (top) and HeLa (bottom) cells. **e.** Western blot analysis of protein extracts from non-transfected HEK293t cells (Control, Con.) compared to cells transfected with SNAP-tdTom (NO stop) or SNAP*tdTom (Stop). Blots were probed with anti-myc (ORF1; top blot) and anti tdTom (ORF2; bottom blot).

To examine whether sequences of ORF1 or ORF2 may have encoded, unintentionally, cryptic proteolytic sites, promoters, ribosome entry sites (or else, e.g.,^62,63^), we replaced ORF1 (i.e., SNAP; 540 bps) by a fluorescent ca^2+^-indicator (GCaMP7s^64^; ∼1400 bps). Importantly, these show no sequence homology. However, this substitution still endorsed functional expression of ORF2 in HEK cells, primary glia, and neurons (**Suppl. Fig. 3a-d**). Similar observations were obtained when ORF2 was replaced by GFP, namely GFP fluorescence could be detected in both HEK and HeLa cells (**Suppl. 3e**). We next examined a much larger ORF1 (a voltage-gated potassium channel Kv4.2^65^; ∼2000 bps) tagged with GFP, when these were separated by a short out-of-frame linker following the stop codon (**Suppl. 4a**). We could easily detect green fluorescence, using a standard electrophysiology set-up, that filled the entire cytoplasm of the cell, showing that it was not bound to channels located within membranes (**Suppl. 4b, c**). This result further rules-out stop-codon readthrough (see above). Together, we find that stop codon-mediated *polycistronicity* is pervasive in a variety of cell types and yields seamless expression of various ORFs. Importantly, translation of ORF2 can be obtained past >2000 bps from the primary initiation site; further extending the limits of *polycistronism* in mammalian cells^66^.

### Kozak sequence enhances stop codons polycistronicity

To determine the constraints of stop-codon mediated polycistronism, we varied the linker bridging ORF1 and ORF2, as well as replaced tdTom by mCherry (**Suppl. 5a)**. We observed polycistronic expression of ORF1 and ORF2, regardless of the preceding sequence (**Suppl. 5b**), although greatest expression of ORF2 was obtained when the linker between the stop codon and ORF2 included a single base-pair causing a frame-shift (**Suppl. 5c, bottom panel, cyan**), potentially resulting from a strengthened *Kozak* sequence^67^. This modification also yielded better expression of ORF1, as noted above (**Suppl. 5c, top panel, cyan**). Conversely, weakest expression level and efficiency were obtained when the methionine was removed from ORF2 (**Suppl. 5c, d**), however this had minimal effect on ORF1’s expression, contrasting the effect of IRES promoters that tend to decrease the expression of ORF1 (**Suppl. 5b-c, no Met, orange**)^32^. Thus, efficient translation of the second (and improvement of the first) ORF is promoted by a *Kozak* sequence (whether canonical or less so) but requires a methionine.

### Development of a genetically-encoded rainbow indicator (GEFRI) for functional highlighting of neurons

To examine whether cells could support *polycistronism* of three ORFs cloned in single file, we introduced three ORFs in the same vector, namely CFP (blue), GCaMP7s (green) and tdTom (red); each ORF starting with a methionine (M) and ending with a stop codon (TGA), and in clone #2, we also introduced a canonical *Kozak* sequences before each M. (**Suppl. 6a, #1, 2, respectively**). All ORFs from both clones expressed well in HEK cells, displaying a large combinatorial diversity of expression patterns, resulting in variations in hues, reminiscent of *Brainbow*^68^ (**Suppl. 6a, micrographs**). We separated the various colors by spectral fingerprinting (**Suppl. 6b**) and assessed expression in individual cells, which demonstrated the improvement in the expression of ORF3 by the *Kozak* seq. (**Suppl. 6c**). Both clones exhibited sufficient expression of ORF2 (i.e., GCaMP7s), attested by the robust reporting on Ca^2+^-activity in HeLa cells (**Suppl. 6d**). We thereby denoted these clones Genetically Encoded Functional Rainbow Indicators (GEFRIs). Similar results were obtained in primary cultured neurons, in which GEFRI_1_ or GEFRI_2_ yielded stochastic expression of the three ORFs (**Fig. 2a, b**). Multicolor fluorescence could be discerned throughout soma, and processes (**Fig. 2b, bottom; purple axon**), from which we could also monitor neural activity (**Fig. 2b, top traces**). Thus, GEFRIs operate as *functional highlighters*^27,69^. To try to examine whether exists any relationship between the expression of the various ORFs, we inverted ORF1 in GEFRI (maintaining length and composition of the DNA) to minimize changes to the clone for better comparison (**Suppl. 7a**); expecting to lose expression of CFP but not of subsequent ORFs. Strikingly, abolishing expression of ORF1 almost completely abolished expression of subsequent ORFs (**Suppl. 7b, c**). To examine the potential relationship between the expression of the first and subsequent ORFs, we produced a stable cell line (from a single clone to minimize variability between cells, see **methods**) that expresses GEFRI_2_ (**Suppl. 8a**). Interestingly, we consistently obtained two main cellular populations (from the seven potential possibilities), namely a population of cells in which we detected expression of ORF1 and ORF3 (encompassing ∼54% of cells) and another population in which we detected all three ORFs (∼41%) (**Suppl. 8b, pink and brown, respectively, summarized in c**). Very few cells showed expression of just a single ORF (e.g., CFP only was observed in ∼4% of cells), and in even fewer cells could we detect exclusive expression of the downstream ORFs (i.e., green and red), consistent with the reversed ORF1 experiment shown above (see **Suppl. 7**). This tight association was further complemented by observing a relatively strong relationship between extent of expression of each ORF in these two populations, with highest association between the first and second ORFs (**Suppl. 8b, summarized in d**). These strongly suggest a translation reinitiation (REI) mechanism, whereby the expression of the second and third ORFs is highly dependent on *a priori* translation of ORF1. This supports our above observations showing complete separation between the translated proteins (and not via read-through, RT; see Fig. 1). We next wondered whether REI would be pervasive *in vivo*, knowing that HEK cells (and cultured cells in general) may exhibit capabilities that may not obtained *in vivo*, especially not by neurons^70^. We thereby produced a AAV1-GEFRI_2_ and, following three weeks after injections into rat brains, extracted the brain and produced brain slices, in which we could easily detect neurons of various colors without immunostaining for the different proteins (**Fig. 2c**). Thus, this clearly demonstrates the *polycistronism* of three ORFs can be obtained *in vivo* (**Fig. 2c**).

**Figure 2.**
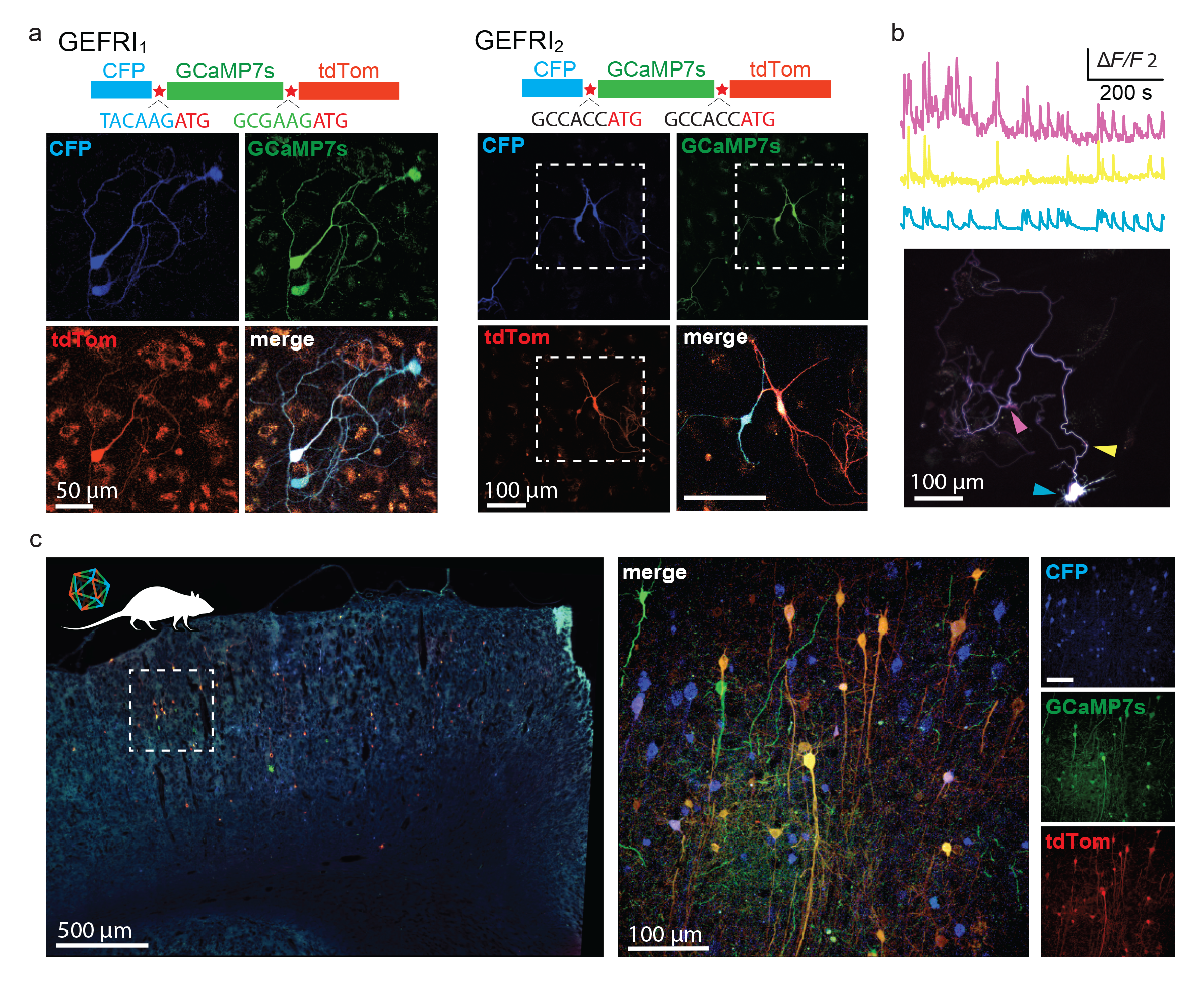
Polycistronic expression of three ORFs in primary neurons *in vitro* and *in vivo*. **a.** Micrographs of primary cultured neurons expressing GEFRI_1_ (left) or GEFRI_2_ (right). Note that whereas all neurons express the first ORF (blue), the expressions of the second (green) and third (red) ORFs are variable, though pervasive. The diverging expression patterns yields variable hues in different neurons. **b.** Representative traces of typical ca^2+^-activity (somatic-cyan and at processes, pink and yellow) from the expression of the middle ORF, GCaMP7s (green), in primary cultured neurons expressing GEFRI_2_. **c.** Micrograph of a fixed brain slice taken from a rats infected with AAV1-GEFRI_2_ three weeks prior. Inset (dashed region)- High power micrograph showing the expression of all three ORFs. Individual channels are shown on the right.

### Harnessing polycistronism in mammalian cells for rapid engineering of novel split-GFP variants

Having observed pervasive *polycistronism in vitro* and *in vivo*, and following our description of the very basic rules of engagement of the process^44^, we proceeded to examine whether *polycistronism* could be employed for splitting proteins with greater ease and rapidity. We initially examined whether we could reproduce the original and prototypical split-GFP (**Suppl. 9a**)^2^. To do so, we simply introduced a stop codon (and methionine, see above) past residue K214 in superfolder-GFP (sfGFP), resulting in two fragments of 214 and 24 residues, namely GFP_1-10_ and GFP_11_-like, respectively (**Suppl. 9a, b; GFP_1*M_**). In HEK cells, we observed moderate expression efficiency (∼25%), with cells showing weak, although specific, green fluorescence (**Suppl. 9c left panels**). This expression did not result from RT as a clone with a frameshift preceding ORF2 equally showed specific, though weaker, GFP-fluorescence (**Suppl. 9c, GFP_1*shift_**). Notably, this fluorescence was necessarily obtained from the expression of both fragments, as the first ORF on its own (isolated GFP_1-10_; in which fragment the chromophore resides) did not yield any fluorescence (**Suppl. 9, isolated GFP_1-10_**). Thus, the original split-GFP could somewhat be obtained by the simple insertion of a stop codon and methionine, however exhibited weak fluorescence and expression efficiency. We deemed this to likely result from instability of one of the fragments, and noted that our 11^th^ β-strand varied quite extensively from the reported one by containing the entire 24 residues, as opposed to a truncated version (of 16 residues) as is found in the original split-GFP^2^; this truncation was indeed necessary for fragment stabilization (**suppl. 9a, see full GFP vs. original GFP_11_**). We then proceeded to try to engineer a new split-GFP variant, which would require less modifications (such as truncations) than the original clone. We introduced a single stop codon in the middle of sfGFP following residue 145; dividing sfGFP into two parts: GFP_1-6_ (six strands; 145 aa) and GFP_7-11_ (five strands; 93 aa), denoted sfGFP_2*_ (**Fig. 3a**). This variant yielded robust expression efficiency (∼90%) with moderate fluorescence (**Fig. 3b**), which was easily improved by the addition of a methionine past the stop codon (**Fig. 3a, b; sfGFP_2*M_**). We then shortened the flexible linker adjoining the two β-strands, denoted split-GFP_2*Del_ and split-GFP_2*MDel_ (**Fig. 3a, green residues**). These attempts yielded reduced expression efficiency, though split-GFP_2*Del_ exhibited very strong fluorescence within the few cells that did show expression (**Fig. 3b**). We then physically separated split-sfGFP_2*_ and split-GFP_2*Del_, by cloning each ORF into independent plasmids (**Fig. 3c**), and observed that both yielded complete reconstitution of GFP, wherein split-sfGFP_2_ exhibiting higher fluorescence (**Fig. 3d, e**). Importantly, expression of the fragments alone did not yield any detectable fluorescence (**suppl. 10a**). To test the quality of complementation of the fragments, we introduced photoactivatable (PA) mutations^27,72^, denoted PA-split-sfGFP_2_ and PA-split-sfGFP_2Del_. Briefly, photo-activatability requires correct alignment and apposition of multiple residues within the GFP barrel for efficient proton transfer^73^. Both variants underwent robust photoactivation following 405 nm illumination bouts (**Fig. 3f, g and h; 55 and 90-fold, respectively**), and only when both fragments were expressed jointly (**Fig. 3f vs. Suppl. 10b**). Interestingly, PA-split-sfGFP_2Del_ underwent more pronounced photoactivation (∼2-fold, **Fig. 3h**). Scrutiny of its sequence revealed that this likely results from the removal of the H149 residue^72^. Regardless, these results show that in both instances the fragments undergo correct complementation, attested by bright fluorescence, as well as reconstitute the internal proton wires and, thereby, photoactivation process. Though beyond the scope of this work, our exploration reveals that the four residues within the linker (F146-H149) are deleterious to the expression and/or fluorescence of split-sfGFP_2Del_, but beneficial for extent of photoactivation (**Fig. 3h**). Thus, our method proved successful for quickly engineering unique split-GFP variants (and split-PA-versions) featuring the entire sequence of GFP compared to *all* previous split-variants^3^. Notably, this splitting point was not predicted by a recent computational approach (**Suppl. 11**)^19^. Therefore, insertion of a single stop codon is a rapid and feasible approach for the quick engineering of split proteins, seamlessly.

**Figure 3.**
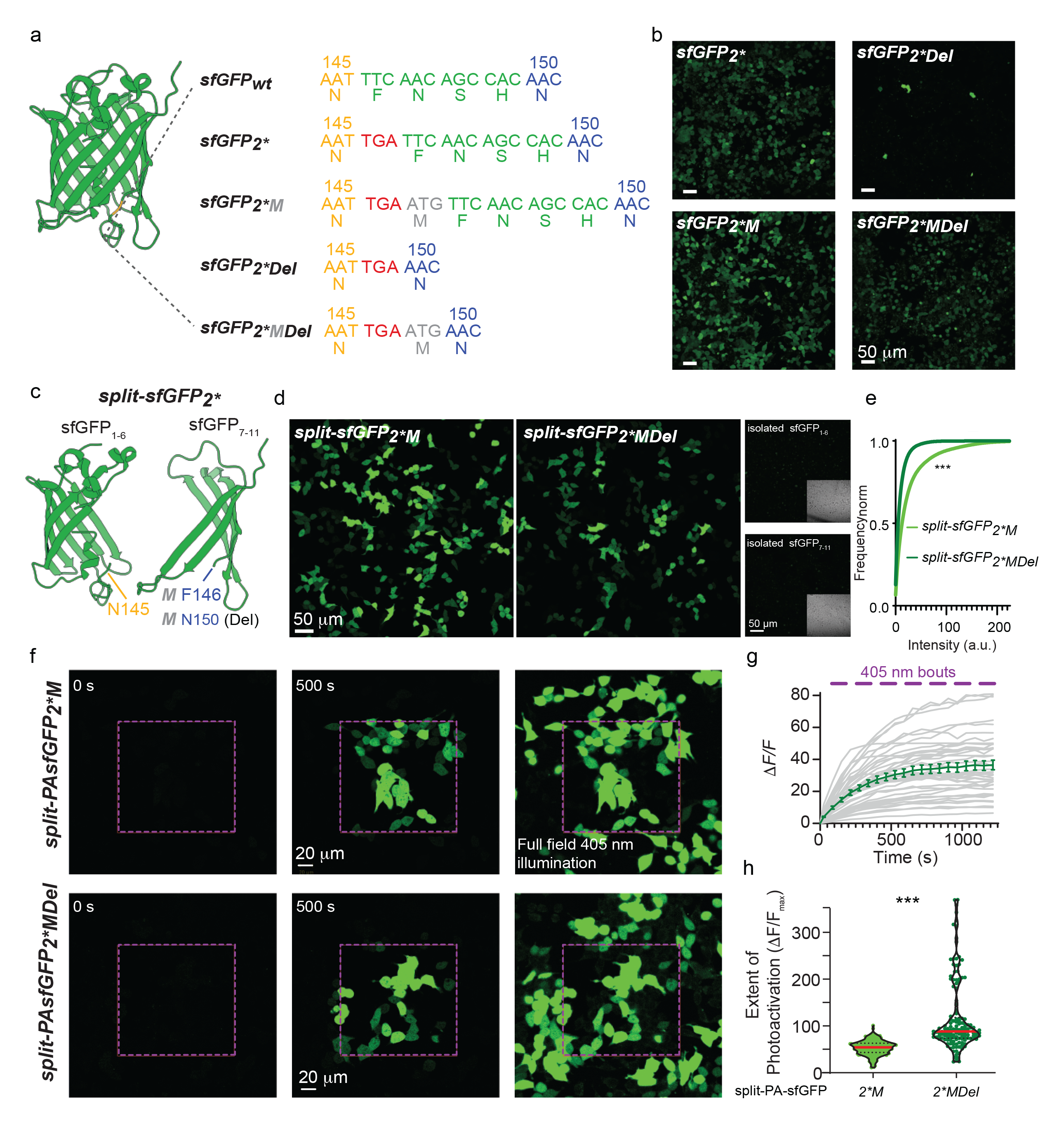
Engineering new split-GFP variants using stop codons. **a.** Illustration of sfGFP (PDB: 2B3P) with the sequence of interest (yellow) into which we introduced a stop codon (TGA, red), following residue N145 (145, yellow). This was thjen followed by introduction of a methionine (grey, *M*) and deletions (*Del*) of residues 146-149 (green residues); yielding four different sf-GFP2* variants. **b.** Micrographs of HEK293 cells expressing the various clones shown in (**a**). **c.** Illustration of the full separation of sfGFP_2\**M*_ and sfGFP_2\**MDel*_ into two independent vectors to obtain two distinct proteins. Separation was performed exactly at site of the stop codon (after N145); isolating the first 6 strands of sfGFP from the 7-11 strands (i.e., sfGFP_1-6_ and sfGFP_7- 11_, respectively). **d.** Micrographs showing fluorescence only when both fragments are co-expressed. in HEK293 cells. Isolated fragments yield no fluorescence (small micrographs and transmitted light insets). **e**. Split-sfGFP_2\**M*_ yields higher expression levels than split-sfGFP_2\**MDel*_. Kolmogorov-Smirnov comparison shows significantly higher expression of split-sgGFP_2_ (***, p<0.001). **f.** Split-PA-sfGFP variants (from **e**) undergo robust photoactivation. Micrographs of HEK293 cells co-expressing both fragments of split-PA-sfGFP*_2*M_* (top) or split-PA-sfGFP*_2*MDel_* (bottom). Micrographs show fluorescence of the clones prior photoactivation (0 s), after photoactivation of a defined region (dashed magenta box) at 500 s, and at the end of the session with full 405 nm illumination of the entire field of view (right), showing all cells that have expressed the clones but were not photoactivated at 500 s. **g.** Time course changes in fluoresce Representative changes of split-PAsfGFP_2_ fluorescence per cell (gray plots) and the average (green; ± SEM), following each photoactivation. **h.** Summary of extent of photoactivation of split-PA-sfGFP*_2*M_* (left) and split-PA-sfGFP*_2*MDel_* (right). Statistics were performed using non-parametric Mann-Whitney test. ***, p<0.001.

### Polycistronism for engineering split-GECI

To try to split more complex probes, in particular GCaMP6s^24^, we rationally introduced stop codons between its three different ORFs (M13; circularly permuted GFP, cpGFP and CaM); expressed the clones in HEKs in which we examined expression (basal fluorescence) and functionality (i.e., Ca^2+^-activity) (**methods and Suppl. 12a**). The clone with a stop codon introduced after M13, specifically following residue E26, provided no basal fluorescence and was not functional (**Suppl. 12a, b, GCaMP6s_1*_**). A stop codon in the middle of the cpGFP yielded a highly functional probe, yet of very low expression efficiency (∼5%; **Suppl. 12a, b, GCaMP6s_2*_**). Lastly, insertion of a stop codon prior CaM (prior residue L268) yielded a probe that expressed well and exhibited robust Ca^2+^-activity (**Suppl. 12a, b, GCaMP6s_3*_;** ΔF/F ∼2). We therefore examined this splitting site, by moving the stop codon within the linker (LP) (**Suppl. 12, GCaMP6s_3.1*, 3.2*_**). In both instances, the performances of the clones were diminished compared to when ORF2 was initiated by leucine (i.e., in GCaMP6s_3*_) (**Suppl. 12b**), consistent with reports showing that leucine is a viable alternative initiation codon^74^. We thereby proceeded to optimize GCaMP6s_3*_ and created a fully split version (M13-cpGFP). In cultured hippocampal neurons, the sole transfection of GCaMP6s_3*_ or M13-cpGFP yielded noticeable basal fluorescence (**Suppl. 12c, d; top panels**), though responses were almost undetectable (**Suppl. 12d, top micrograph and traces**). Greater responses were obtained when the soluble CaM (MLP-CaM) was co-transfected with the first fragment (**Suppl. 12c, d, bottom micrographs and traces, summary in e**). Advantageously, these demonstrate that the endogenous CaM molecules in neurons did not serve as functional partners for the M13-cpGFP fragment (**Suppl. 12e**), as previously demonstrated to be a nuisance when using other calcium probes (e.g, ^75–77)^. Electrophysiological recordings coupled with fluorescence imaging of neurons co-transfected with M13-cpGFP and MLP-CaM showed that the split probe maintained some of the key features of the parent probe, namely reports on single action potentials^24^ with comparable kinetics, with larger responses at processes than soma as seen with other probes^27,78^ (**Suppl. 12f, g**). To be relevant *in vivo*, we proceeded to test whether viral infection would produce sufficient expression of split-GCaMP6s_3_, as viral infections typically result in lower protein expression levels compared to chemical transfection methods (see **methods**)^79^. We thereby produced two AAVs, each packaged with a different fragment of split-GCaMP6s_3_, and infected cultured primary hippocampal neurons. Fluorescence could be easily detected in most neurons that were solely infected with the M13-cpGFP-virus (**Suppl. 12h, top and bottom panels)**, in which very weak Ca^2+^-activity could be monitored (**Suppl. 12h, top and bottom traces, dashed regions**), consistent with our above results (see **Suppl. 12c**). Importantly, dually infected neurons exhibited ∼10-fold larger responses in both soma and processes of neurons (**Suppl. 12i, top and bottom** panels and traces; summary in j).

### Optimizing split-GCaMP6s

The observation that a sole fragment of the split-GCaMP6s_3*_ was both fluorescent and active (even if modestly, see **Suppl. 12**) motivated us to try to engineer a different probe that would necessarily be fluorescent and ca^2+^-responsive—solely after complementation. We reverted to re-examine GCaMP6s_2*_ as this probe exhibited lower efficiency (i.e., expressed only in a handful of cells), albeit showed bright fluorescence and activity in a handful of cells, suggesting it is less stable (**Suppl. 12b**). We thereby initially tried to improve its stability by modifying its cpGFP by introducing superfolder (sf)^27,80^ or optimal-sf (osf)^4^ mutations, denoted sfGCAMP6s_2*_ and osfGCAMP6s_2*_, respectively (**Suppl. 13a, stars**). Indeed, this optimization step yielded variants with considerably higher expression efficiency, with sfGCaMP_2*_ exhibiting significantly larger calcium responses compared to osfGCaMP6s_2*_ (**Suppl. 13b, c**). We then separated the clone into two plasmids and found that fluorescence and activity were exclusively obtained when both fragments were co-expressed (**Suppl. 13b vs. d**). We thereby denoted this clone split-GCaMP6s_2_. Interestingly, the activity of split-GCaMP6s_2_ in HEK cells was similar whether the reconstituted probe was obtained from the stop-codon containing plasmid (i.e., sfGCaMP6s_2_*) or from the co-expression of the two independent plasmids (**Suppl. 13a, b, top and bottom lanes**). This further demonstrates the power of the approach to obtain relevant expression levels of the post-stop codon ORF specifically in HEK cells. Split-GCaMP6s_2_ expressed well and was functional *in vitro* in cultured neurons (readily reporting on single action potentials as the parent probe) (**Fig. 4a-c**), as well as permitted prolonged imaging of activity *in vivo* in larval Zebrafish (**Fig. 4d**). Lastly, akin to split-sfGFP, this clone also exhibited proper complementation, assessed by the introduction of the PA mutations, denoted split-PA-sfGCaMP6s_2_ (**Fig. 4e-g**). To examine whether this splitting site would be suitable for other GECIs, we introduced a stop codon in the middle of the fluorescent protein of the red jR-GECO1a indicator^81^, specifically within the cp-mApple following residue S120 (**Suppl. 14a, b**), and find it to be functional. Co-expression of the fully separated jR-GECO1a (split-jR-GECO1a) yielded robust expression and ca^2+^-responses in HHEK cells (**Suppl. 14, jR-GECO1a_*120_**), whereas each fragment on its own did not show any basal fluorescence nor detectable activities (**Suppl. 14b, bottom**).

**Figure 4.**
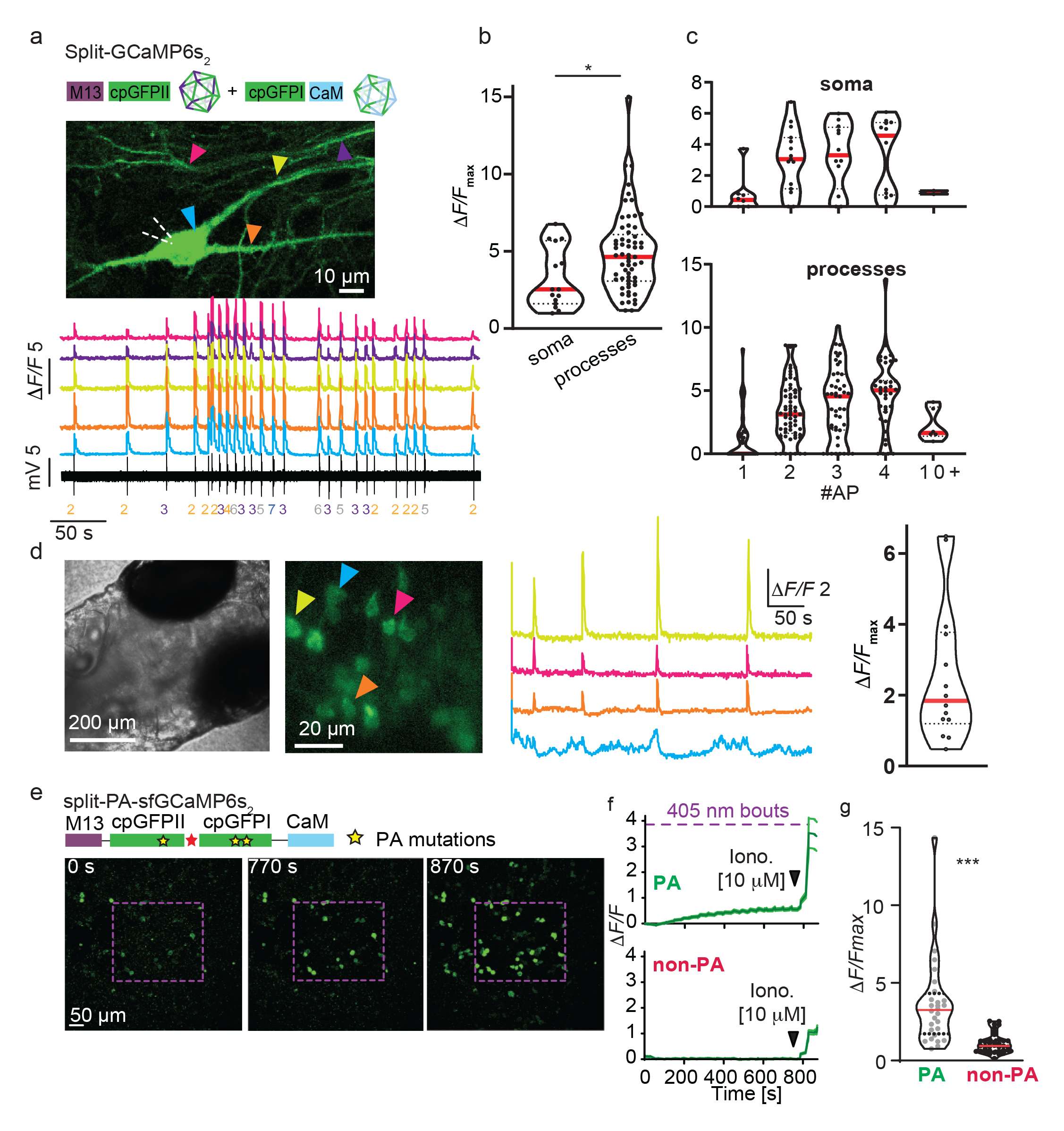
Split-GCaMP6s_2_ reports on neuronal activity *in vitro* and *in vivo*. **a.** Micrograph of a pyramidal neuron co-infected with two AAVs encoding for the two fragments of split**-** GCaMP6s_2_ (cartoon). The neuron is simultaneously loose-patched (dashed white lines highlights pipette). (below) Color-coded fluorescent traces (corresponding to regions noted in micrograph) that correspond to neuronal firing. Number of action potentials (#APs) are noted below patch recording (black trace). **b.** Summary of maximal Δ*F/F* obtained in soma and processes, and correspondence towards #APs (**c**) (n = 11). **d.** Micrographs of zebrafish larva expressing both fragments of split**-**GCaMP6s_2_ (see **methods**), and traces of spontaneous ca^2+^-activities from select neurons highlighted in fluorescent micrograph (color-coded arrowheads); summarized in violin plot (right). **e.** Engineering a photoactivatable superfolder split-GCaMP6s_2_ (split-PA-sfGCaMP6s_2_). Cartoon representation of the location of PA mutations (yellow stars) within the domains of the clone. Micrographs of HEK cells, expressing split-PA-sfGCaMP6s_2_ (with a stop codon, top cartoon), before (0 s) and at the end of photoactivation sessions consisting of 405 nm bouts of illumination within dashed magenta region. At the end of the session, all cells were exposed to 10 μM ionomycin to saturate all cells Ca^2+^. **f.** Summary of photoactivation and ca^2+^- responses of split-PA-sfGCaMP6s_2_. Traces show responses (Δ*F/F*) of cells located within the region of interest (dashed region as in **e**, i.e., photoactivated cells, PA in green) (top panel) or cells outside the region photoactivation (non-photoactivated, non-PA in red) (bottom panel). Magenta bar represents bouts of targeted 405 nm illumination. **g.** PA cells exhibit larger responses to ca^2+^ than non-PA cells. Summary of the maximal responses (Δ*F/F_max_*) of photoactivated and non-photoactivated cells to ionomycin.

### Engineering of split-MEGIC-a ca^2+^-reporter localized at mitochondria-ER contact site

To benefit from the unique features of a split-GECI, we sought to design a probe that would express at two different ca^2+^-handling organelles that come into very close proximity within cells, such as mitochondria-ER contact sites^82–84^. We first examined whether our split-GFP variants could be reconstituted at these contact sites. For this, we tethered the GFP_1-6_ fragment to TOM20 (a mitochondrial resident protein^85^), and the GFP_7-11_ fragment to the transmembrane domain of the ER resident protein CB5^86^, to which we have also included mRuby3 at the carboxy terminus for visualization of expression (**Suppl. 15a, Part I and II, respectively**). In HEK cells, as its parent split-GFP_2_, neither of the clones yielded any detectable fluorescence when expressed individually, even though mRuby3 could be observed in ER membranes that expectedly delineated the nucleus (**Suppl. 15b**). Co-expression of both parts yielded green punctuates that co-localized with red fluorescence from the ER (**Suppl. 15c**). To validate localization of Part I onto the mitochondrial surface, we triple co-expressed part I, a soluble GFP_7-11_ fragment and mito-CAR-GECO1 (a red GECI localized to mitochondria^87^). This combination surface labeled (green) all mitochondria (red) (**Suppl. 15d**).

We next replaced the GFP fragments with those of split-GaMP6s_3_, yielding a probe we denoted MEGIC— <u>M</u>ito-<u>E</u>R <u>G</u>enetically Encoded <u>I</u>ndicator for <u>C</u>a^2+^. We expressed the first part of MEGIC (consisting of a mitochondria localized M13-cpGFP fragment) and noted that in HEK cells it exhibited basal fluorescence (i.e., prior ca^2+^-activity), which was restricted to mitochondria, as expected (**Suppl. 16a**). Unexpectedly, the first fragment on its own (i.e., only half of the probe) was functional in HEK cells, in which could see matching activities between the MEGIC fragment and the inner mitochondrial red probe mitoCAR-GECO^87^ (**Suppl. 16b**). However, co-expression with the second fragment significantly improved the amplitude of the responses >2 fold (**Suppl. 16c, d**). A similar picture emerged when MEGIC was expressed by viral infection in cultured neurons. Specifically, green fluorescence could be observed throughout neuronal somata and processes when only the first part was expressed (**Suppl. 17a**), unlike its parent probe split-GCaMP6s_3_ (see **Suppl. 12**). Co-expression of both fragments significantly improved the responses to Ca^2+^ (**Suppl. 17b, summary in c**). Thus, the tethered variants can employ the endogenous CaM as partner for activity, whereas the soluble fragment does not^88^. Interestingly, the responses of the probe were highly diverse, depicting ca^2+^-events ranging from seconds to minutes (**Suppl. 17d, e**) and these did not match with action potential firing, unless the neurons fired high frequency bursts (**Suppl. 17f**). Lastly, we imaged MEGIC in vivo, in larval zebrafish, in which we could observe green puncta within red-filled neurons, which readily reported on highly localized spontaneous activities (**Suppl. 17g**).

To try to produce a better probe, we engineered another MEGIC variant (MEGIC2) based on split-sfGaMP6s_2_ (**Fig. 5a**). We observed no fluorescence when each fragment was independently expressed in HEK cells nor in neurons (**Suppl. 18a-d**). The expression pattern differed from MEGIC_1_ (which labeled all mitochondria) and exhibited a unique pattern of expression of fluorescent green puncta (packets) in HEK cells and cultured neurons (**Fig. 5a**). In neurons, most of the green packets were localized juxtanuclear at the soma, with multiple packets distributed along dendrites (**Suppl. 18e and Fig. 5a, b**). Importantly, co-expression of both fragments yielded low basal green fluorescence which increased in the presence of calcium in neurons (**Suppl. 18e and Fig. 5c**). Dendritic and somatic packets sites were very active (without any changes in RFP; **Fig. 5c, red channel and traces**), and many additional sites (especially along processes) were noticeable only during activity owing to very low basal fluorescence (**Fig. 5c**). This feature yielded very high signal to noise; important for imaging brief events within small regions (below). Akin to other calcium probes, changes in fluorescence (i.e., Δ*F/F*) were consistently larger at processes compared to somata (**Fig. 5d**)^27,28^.

**Figure 5.**
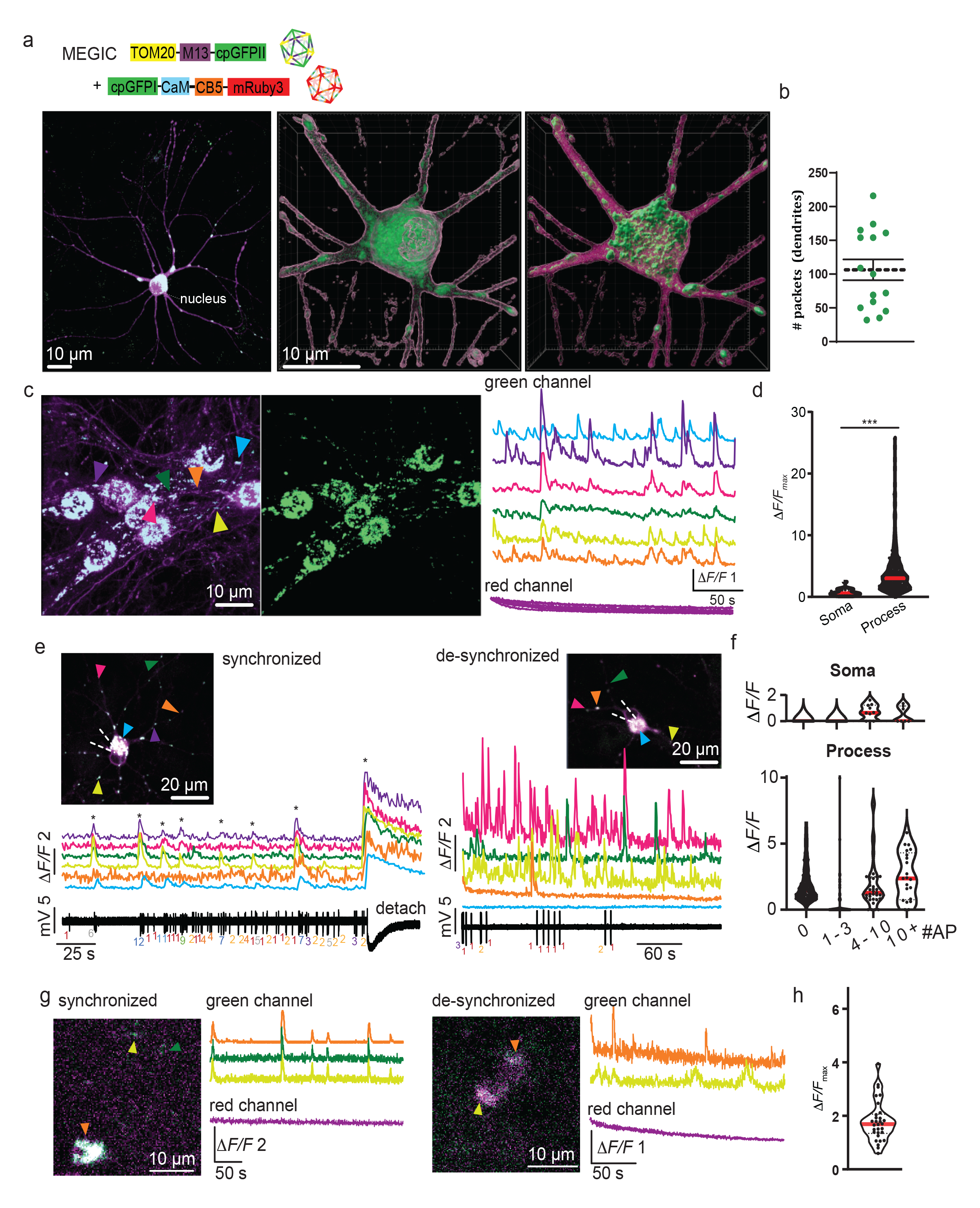
Split-MEGIC in soma and dendrites of neurons, *in vitro* and *in vivo*. **a.** (top) cartoon representation of two independent plasmids employed to produce two separate AAVs (colored icosahedron); (bottom) micrographs showing cultured neurons infected by the two viruses. Magenta highlights ER, whereas green (white; overlap of green/magenta) highlights Mito-ER contact sites (packets). Packets are distributed across the entire cell, with the majority at the soma, juxtanuclear (arrow). Middle and right panels show 3D-reconstructionn of high-power micrographs (pseudo-super resolution) of soma and proximal dendrites showing a large abundance of amorphous packets within the soma, as well as in dendrites, summarized in (**b**). **c.** A micrograph of a group of neurons expressing split-MEGIC, taken during peak activity (right traces, green channel) during which many more packets become noticeable (otherwise completely non-detectable). Red fluorescence does not show any changes in fluorescence (bottom magenta traces). **d.** Summary of Δ*F/F*_max_ in somata and processes. **e.** Representative examples of neurons dually patched (loose patch) and imaged simultaneously (pipette marked by dashed white lines). Ca^2+^- activities at various regions (arrowheads) are shown below micrograph (color-coded). Traces show synchronized activity between multiple sites along the dendrite, including soma, in the same neuron. Synchronized activities (asterisks) corresponded to bursts of action potentials (black trace, # action potentials are noted below each transient). To saturate all sites simultaneously, patch pipette was gently detached from cell. Right micrograph and traces show a different neuron in which many sites along the dendritic tree exhibited variable activity patterns (de-synchronized). These ca^2+^-activities did not correspond to action potential firing (bottom black trace). **f.** Violin plots of Δ*F/F* at soma (top) and processes (bottom) as a function of the number of action potentials recorded simultaneously by loose patch. Note the very low changes observed at the soma (which exhibits very high basal fluorescence). **g-h.** Synchronized and de-synchronized activities *in vivo*, in larval Zebrafish, summarized in (**h**) (N= 3). No changes in red fluorescence were observed (magenta traces), also showing lack of movement in the z-axis.

We observed two different modes of activity, namely synchronous and de-synchronous activity between different mito-ER contact sites within a given neuron (**Fig. 5e**). Synchronized activity matched very high neuronal activity consisting of bursts of action potentials (**Fig. 5e, left**), whereas lower activity (of few action potentials), did not engender any responses (**Fig. 5e, right and summary in f**). This demonstrates that synchronized activity between multiple sites jointly, requires the neuron to be flooded by calcium, which we confirmed by slightly detaching the patch pipette from the cell leading to concomitant responses by all sites monitored (**Fig. 5e, left, detach**). De-synchronized activity did not report on lower firing rates, rather was quite independent of electrical activity, reflecting highly localized ca^2+^-activity within individual contact sites (**Fig. 5e, right and Suppl. 18e**). Similar observations were obtained *in vivo*, in larval zebrafish expressing both independent fragments, in which we could detect change in the fluorescence of the green puncta, without any changes in the fluorescence of the ER-resident RFP (**Fig. 5g, h**).

Interestingly, in aged neuronal cultures (<2-3 weeks), we noted red fluorescence within dendritic spines, depicting presence of ER (**Fig. 6a, magenta**). However, a handful of spines also included green fluorescence (**Fig. 6a, green**). We observed green fluorescence invading and retracting from spines, as well as could detect highly localized calcium activity without changes in the intensity of the red fluorescence (Fig. 6x). In another series of experiments, mitochondria-ER contact sites displayed synchronized and de-synchronized activity among several contact sites along the dendrite. Synchronized activity reflected back-propagating action potentials (likely bursts, see **Fig. 5**) that simultaneously flooded both spines and dendrites, whereas de-synchronized activity was also observed intermittently (**Fig. 6c, right traces, black arrowheads indicate de-synchronized activity**).

**Figure 6.**
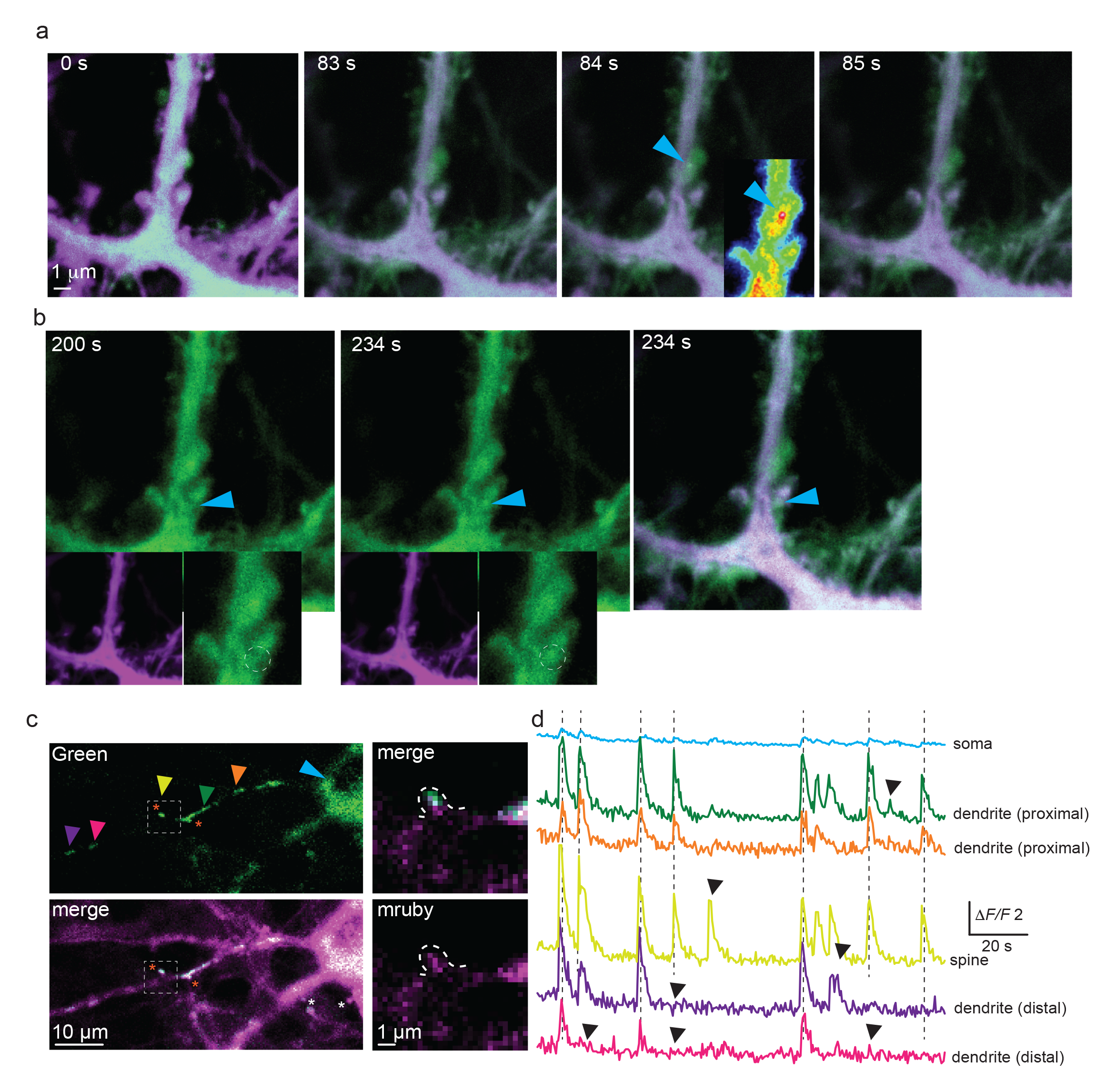
Mito-ER contact sites in dendritic spines. **a.** Sequential micrographs of a dendritic segment from cultured neurons, expressing split-MEGIC2. Ca^2+^-activity is observed in a spine (cyan arrowhead). Spine head is mostly filled with green fluorescence. Activity is highlighted at 84 s (inset-heatmap image showing localized activity in the middle of the spine head). **b.** Growth of spine and filling with a Mito-ER contact site (cyan arrowhead). ER (magenta) can be seen at shaft. Bottom micrographs show corresponding images of the ER (magenta) and high power images of the growing spine (cyan arrowhead and dashed white region). **c.** Synchronized activity between a Mito-ER contact site localized within a spine head. Micrographs shows multiple mito-ER contact sites along dendritic shaft, within spines (red asterisks) and soma. ER is seen throughout soma, and dendrites, as well as in several spines (red and white asterisks). Synchronized activity is detected between soma (cyan arrowhead), proximal and distal dendrites and within a single spine (yellow arrowhead). **d.** Synchronized and de-synchronized activity between spines, dendrites, and soma. Traces (colors-coded as in arrowheads in **c**) show highly synchronized ca^2+^-activity between soma (cyan) and several sites along processes (vertical dashed lines. Localized and de-synchronized activity is also observed (black arrowheads).

Lastly, we injected (two independent viruses) split-MEGIC_2_ to brains of mice and imaged activity, *in vivo*, via two-photon (2P) excitation microscopy at multiple sessions (see **methods**)^89^. As in cultured cells, red fluorescence was seen throughout the neurons, whereas green fluorescence was restricted to small puncta in somata and processes of cortical pyramidal cells (**Fig. 7a**). Neurons exhibited a large variety of Ca^2+^-dynamics, ranging from sharp to very prolonged and flickering ca^2+^-activities, not to mention synchronized and de-synchronized among numerous cells, and these dynamics varied from session to session (**Fig. 7b**). Together, these demonstrate that split-MEGIC_2_ is compatible with 2P-imaging *in vivo*. Secondly, our results show the ability of the probe to report on a plethora of complex calcium activities at mito-ER sites within neurons *in vivo*.

**Figure 7.**
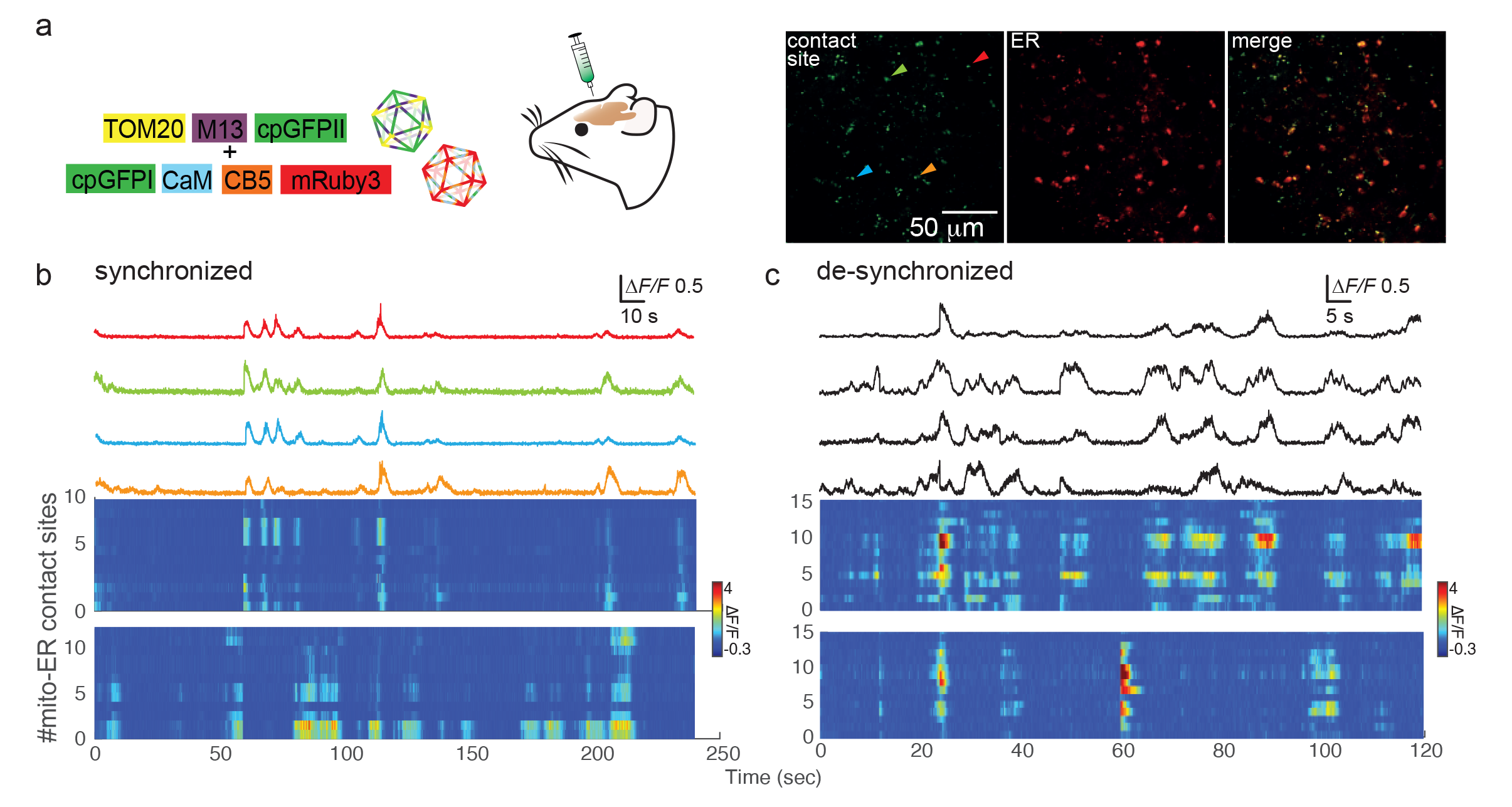
Expression and activity of split-MEGIC_2_ *in vivo* in brains of mice. **a.** Cartoon representation of two independent plasmids employed to produce two separate AAVs (colored icosahedron) that were injected *in vivo*. (right) Micrographs showing infected neurons expressing split-MEGIC2 (red, ER; green, mito-ER contact sites). Images were acquired at 300 µm below pia by a two-photon excitation microscopy. Arrowheads depict regions for which activities are shown in corresponding colored-coded traces in (**b**). **b.** Representative traces depicting highlysynchronized Ca^2+^-activities in multiple sites. Traces are colored as arrowheads in (**a**). (below) Raster plots from two independent sessions showing recordings from a ∼dozens mito-ER contact sites randomly selected. **c.** Traces and raster plots from two other sessions in which we observed different patterns of ca^2+^-activities, many of which were de-synchronized between various Mito-ER contact sites.

## Discussion

We demonstrate non-canonical polycistronic expression of up to three ORFs cloned in-sequence, and solely separated by stop codons, *in vitro* and *in vivo* (e.g., ∼1400 bps- **Suppl. 3,** and ∼2000 bps- **Suppl. 4**). These results deviate from current understandings whereby non-canonical post-stop codon translation, such as by REI or LRS^39–41^, are thought to ensue primarily following significantly shorter distances following the primary AUG or 5’ UTRs (e.g., 100-400 bps)^44,55,90^. Longer distances have been less explored, as these mechanims are typically observed in instances of premature stop-codons occurring early in ORFs^43,49^. Moreover, the few reports that do describe eukrayotic polycistronic translation following longer ORFs typically involve cryptic promoters or IRES sequences^91,92^, whereas translation via REI or LRS (e.g., ^93^) is more rare or often mistaken for RT^94^. Importantly, and to the best of our knowledge, the rules and propensity for non-canonical translation of three ORFS have not been demonstrated prior this report, especially *in vivo* as we show (**Fig. 2c**). Thus, here we extend the permissive distances for the occurrence of stop-codon mediated non-canonical translation of up to three different ORFs^61,66^, without excessive add-ons (e.g., enhancers and promoters^37^, IRES^32^, self-cleavable peptides^33,34^) (**Suppl. 5**), and rule-out RT as a possible mechanism (e.g., **Fig. 1a, e**).

Our simple strategy of insertion of stop codons firstly yielded the engineering of a Brainbow^68^-like probe, denoted GEFRI. Akin to Brainbow, GEFRI includes three different ORFs encoding for three different fluorescent proteins (FPs) with unique emission spectra (blue, green, red) (**Fig. 2**). However, in contrast to Brainbow, here the pattern of expression of the various FPs and emission intensity (specifically of the green emitting probe; GCaMP6) do not remain constant (**Fig. 2 and Suppl. 6**). For instance, expression levels of each FP, and hence emerging color within individual cells, is bound to change during different cell states, such as cell division, and green fluorescence of GCaMP6 varies with Ca^2+^-activity, such as action potential firing in neurons (**Fig. 2b, Suppl. 3c, Suppl. 6d**). This probe thereby opens new means to optically probe cells during different states, including division and cellular activity. Though beyond the scope of this work, we also noted that expression stoichiometry can vary quite extensively following different cellular stressors (e.g., **Suppl. 19**).

Another direct output of our strategy was the rapid development of split-GECIs by firsty screening stop codon-based probes specifically in HEK293t cells (see **methods**). We find that HEK293 are ideal for this purpose owing to their tendency to yield large expression levels of proteins, as well as their propensity for non-canonical translations^95^. We exploited these features to obtain sufficiently high expression levels of various ORFs located downstream of stop codons, to hone-in on viable splitting sites. We thereby denote our strategy SPLIT-IT—<u>S</u>top-codon mediated <u>P</u>olycistronic <u>I</u>nduction in TransIently <u>Transfected</u> HEK cells. SPLIT-IT diverges from previous strategies developed, for instance an intein-based approach (that introduces at least six exogenic bps at the splitting site, which may interfere with protein structure and function^21,96^), as it enables to maintain the entire sequence of the protein—seamlessly— without excessive sequences, not even a methionine or Kozak sequences (though the latter greatly improve expression). SPLIT*-*IT enabled the rapid engineering of a palette of novel split-GFP variants (**Fig. 3 and Suppls. 9-11**). More specifically, insertion of a single stop codon within sfGFP, at the original site of splitting the split-GFP (residue K214), reconstituted the original split-GFP variant, although it exhibited very poor performance (**Suppl. 9**). The weak fluorescence and very low expression efficiency suggested to us that the second fragment (past the stop codon) was highly unstable. Indeed, this proved to be true, as reported in the original report (the 11^th^ strand had to be truncated by 8 residues) (**Suppl. 9a, full GFP vs. original GFP_11_**)^2^. We thereby employed SPLIT-IT to find other viable splitting sites. Intuitively, we rationally introduced a stop codon in the middle of sfGFP, following the six^th^ strand (residue 145), resulting in two ∼equal parts: GFP_1-6_ (145 aa) and GFP_7-11_ (five strands; 93 aa)—which yielded a new and highly functional split-GFP variant. Expectedly, addition of methionine improved this clone’s expression. (**Fig. 3a, b; sfGFP_2*M_ and sfGFP_2*_, respectively**). We then tried to follow the methodology of previous reports and truncated the flexible linker between the six^th^ and the seven^th^ strands by five aa (see **Fig. 3a**), although this resulted in the drastic reduction in expression efficiency. (**Fig. 3a, b; sfGFP_2*MDel_ and sfGFP_2*Del_, respectively**). This suggested that the truncated residues (flexible linker) are essential for the quality of the GFPs fluorophore reconstruction. Indeed, we obtained better expression of the fully separated and non-truncated split-sfGFP_2_ (**Fig. 3c, d, e; split-sfGFP_2_ vs. split-sfGFP_2Del_**). These strongly argue that insertion of a single stop codon in a protein’s sequence can predict the quality of the split-proteins functionality. We further introduced PA mutations^27,72^ to test the quality of complementation of split-GFP_2_ variants. While both PA-variants underwent photoactivation following 405 nm illumination bouts, on par with values obtained for the regular PA-GFP^27,72^ (**Fig. 3f, g and h**), PA-split-sfGFP_2Del_ underwent a more pronounced photoactivation (∼2-fold higher than split-sfGFP2, **Fig. 3h**). Scrutiny of its sequence revealed that this likely resulted from the removal of the H149 residue^72^, however removal of four residues within the linker (F146-H149) are deleterious to its expression. Thus, our method proved feasible for splitting GFP, but also proved valuable for discovering residues involved in the photoactivation process of GFP molecules. Together, SPLIT*-*IT proved successful for generating a new—and seamless—functional split-GFP featuring its entire sequence. Interestingly, this site was not predicted by a recent computational approach (**Suppl. 11**).

We exmained SPLIT-IT for generating self-complementing split-GECIs. We tested a handful of splitting sites, with the aim to invest as little effort as possible in their engineering, so as to obtain viable (even if not the most oprimized) clones rapidly (**Suppl. 12-14**). These clones, unlike previous split proteins^18^, did not require excessive optimization as typically done. We rationaly introduced stop codons prior or within the linkers of GCaMP6s^24^ (**Suppl. 12**) to minimally interrupt the folding of the protein’s domains^97^. Simple insertion of a stop codon following the cpGFP (prior 268L) yielded a clone that expressed well and exhibited robust calcium activity (**GCaMP6s_3*_**; **Suppl. 12a**). Notably, ORF1 on its own, namely the M13-cpGFP fragment, was detectable and functional on its lonesome, yet its performance was drastically improved when co-expressed with the soluble CaM fragment (**Suppl. 12d-j**). Further, split-GCaMP6s_3_ successfully reports on single action potentials (**Suppl. 12f, g**), showing it retained the sensitivity of its parent GCaMP6s. These observations imply the possibility of using M13-cpGFP fragment as a calcium reporter that can be used to indicate the extent of expression of CaM molecules in different regions of the cells, such as synapses^99^. Moreover, this probe may be less disruptive to ca^2+^-dependent process, by providing no additional ca^2+^-buffers to cells. Notably, substitution within the linker between the first fragment and CaM (i.e., residues 268L or 269P in GCaMP_3.1*_ and GCaMP_3.2*_, respectively) drastically decreased its ability to respond to Ca^2+^ elevations (**Suppl. 12a, b**), supporting previous reports showing the importance of this linker in this probe^100–103^, but also fluorescent probes (e.g., ^104^).

Interestingly, insertion of a stop codon in the middle of the cpGFP (GCaMP6s_2*_) yielded probes of very low basal fluorescence and large responses, but of very poor expression efficiency (∼5%; **Suppl. 12a**). The low expression efficiency, along the high functionality of GCaMP6s_2*_, strongly suggested instability of the fragments, which thereby yields very low amounts of functional probes in cells. To address this, we introduced various types of superfolder-mutations^27,80^ which enhanced efficiency and functionality more than four-fold (sfGCAMP_2*_; **Suppl. 13**). Following physical separation of the fragments (i.e., split-sfGCAMP_2_), the probe maintained its high functionality (*ΔF*/*F*∼11; **Suppl. 13**), reporting on locale Ca^2+^-elevation in processes, and on single action potentials *in vitro*, in cultured primary neurons (**Fig. 4a-c**), as well as *in vivo* (**Fig. 4h**). Importantly, in opposite to the “leak” fluorescence and activity observed in other clones, (e.g., GCaMP6s_3_; **Suppl. 12**), this probe was only fluorescent and active when both fragments were co-expressed (**Suppl. 13 c-d**). We further introduced mutations that enable photoactivation, as in our PA-split-GFP variants, leading to the generation of functional split-PA-sfGCaMP6s_2_ (**Fig. 4**). This is consistent with previous reports showing the benefits of superfolder-mutations towards expression of photoactivatable probes at temperatures above 34° C^27,30^. Lastly, this splitting site was found to be compatible with a red-GECI (jR-GECO1a^81^), specifically when the latter was separated at residue S120, and with its entire sequence intact (**Suppl. 9**). These suggest that this splitting site could be generalized towards other GFP/RFP-based probes, which can strongly argue on the globality of this splitting site for other GECIs. Together, our observations demonstrate the reliability of the SPLIT-IT approach and, importantly, its simplicity.

Having obtained the desired clone, namely that exhibits no fluorescence or activity prior complementation, we then leveraged it for reporting on ER-mitochondria Ca^+2^ dynamics, thereby denoted split-MEGICs (**Fig. 4, Suppl. 16 and 17**). Split-MEGIC probes encompass the advantages of ER-mito FRET or BRET-based activity probes and of split-GFP markers that highlight these regions (^105–107^, and see review ^108^). Analogous to our observations above, we found that part1 of split-MEGIC_3_ (i.e., TOM20-M13-cpGFP which is based on split-GCaMP6s_3_) can interact with the endogenous calmodulin and yield fluorescence and responses, whereas co-expression of both fragments significantly improved the responses to Ca^2+^ (**Suppl. 16, 17**). However, diverges from its parent, as its responses are significantly higher. Nevertheless, this probe overestimates the number of contacts as it yields fluorescence on top of all mitochondria that are not necessarily in contact with the ER (**Suppl. 15**). These results raise the possibilities that the apparent concentration of CaM may be higher around mitochondria (for instance, by its immobilization onto other proteins^109^) or, and perhaps more probable, that mito-ER sites are of higher Ca^2+^ concentrations.

To bypass the limitations of plit-GECI_1_, we produced another MEGIC variant, based on split-GCaMP6s_2_, which was fluorescent and functional only when both fragments were co-expressed (**Suppl. 18a**). Thus, split-MEGIC_2_ is a more reliable ER-mitochondrial probe^108^. We observed hundreds of puncta per HEK cell (although of much lower density than the total number of mitochondria, see **Suppl. 15**), most of which were juxtanuclear, highly consistent with previous observations^110^. Interestingly, in neurons we similarly observed packets of ER-Mito contact sites juxtanuclear at the soma, however multiple sites along dendrites could also be detected (**Fig. 5**). Packets that could be detected prior activity (i.e., not firing bursts of action potentials) (**Fig 5f**) suggests higher calcium levels at these sites, whereas many additional sites could be observed transiently along the dendrite during various types of activities. One interesting site at which we observed a contact site was at dendritic spines (**Fig. 6**). We observed contact sites within red-filled spines, demonstrating both invasion of the ER and mitochondria into the structures, as well as observed green spines in which ER was not apparent, and these tended to be much larger (**Fig. 6**)^111^. Lastly, we could also see a dynamic invasion of green into a spine (**Fig. 6b, arrowhead**). In these and a handful of additional spines, we could monitor very localized activity (**Fig. 6a, cyan arrowhead**), but also synchronized activity between the spine and mito-ER contact sites along the dendrite (**Fig. 6c, light green arrow**). This suggests the occurrence of a burst of back action potentials (bAP), among which the spine also exhibited isolated patterns of activity (**Fig. 6c, traces and black arrowheads**). We would like to emphasize that these observations were very surprising to us, as the occurrence of ER in spines is relatively low^112^, and even more rare are spine-mitochondria ^111,113–115^. To the best of our knowledge—the occurrence of an active mito-ER contact site within an excitatory synapses (i.e., spine) has never been previously reported^111,115,116^.

Together, we provide user with a unique means to develop split protein, as well as engineered a palette of split probes, notably Split-MEGIC_2_. Importantly, we show that the probe can be employed in various cell types to detect various ca^2+^-related at unique sites of the cells, such as spines, *in vitro* and *in vivo* in Zebrafish larva (**Suppl. 17h and Fig. 5**) and mice brains (**Fig. 7**). The latter was particularly useful to demonstrate that the probe is readily suitable for 2PE-microscopy. Together, split-MEGIC_2_ can monitor global, as well as local ca^2+^-elevations at the ER-mito junction. Its immobilization affords the probe excellent spatial resolution, as it can report on local changes in calcium between two adjacent sites (**Fig. 4d**), similar to other immobilized probes (e.g.,^25,117^). We anticipate that split-MEGICs will serve as a steppingstone for the development of additional activity probes for other organelles.

## Materials and Methods

### Mutagenesis and cloning

#### Membrane localized SNAP/CLIP-tags

The SNAP and CLIP-tag mutant (pU6-SNAP and pU6- CLIP) were kindly provided by Prof. Jakobs S, University of Gottingen, as reported^118^. To create pCMV(MinDis)-SNAP and pCMV(MinDis)-CLIP, we used the pCMV(MinDis)-iGluSnFR (Plasmid #41732 Addgene) backbone, which contains a signal peptide and the PDGFR transmembrane domain. The coding sequences of the tags were amplified by PCR reactions using primers: Forward: TATATA**AGATCT**ATGGACAAAGACTGCG and Reverse: TATA**CTGCAG**GCCCAGCCCAGGCTTGC; while inserting *BglII* and *PstI* restriction sites, respectively. We double digested the products and replaced iGluSnFR by either SNAP or CLIP proteins. The FPs td-tomato-C1 (plasmid #54653 Addgene) and eGFP (kindly provided by Prof. Aronheim A, Technion) were inserted by PCR amplification while inserting *XhoI* restriction sites using the primers: forward 5′-TATA**CTCGAG**CCTACTCAGACAATGCGATGCAATTTCCTCAT and reverse 5′- TATA**CTCGAG**GATCATCATCAGGATCATCATCAATGGTGAGCAAGGGCGAGGA. *XhoI* digestion and insertion of the tags into pCMV(MinDis)-SNAP and pCMV(MinDis)-CLIP constructs, resulted with a 12 aa linker between the TGA stop codon at the 3’end of the PDGFR and the 5’end of the FPs (SNAP*tdTom; **Fig. 1b**, and CLIP*GFP; **Suppl. 3e**, respectively). For replacing the TGA (stop codon) at the C-terminus of the PDGFR with Ser aa (TCC), a point-mutation PCR was performed using the following primers: 5’- CTTTGGCAGAAGAAGCCACGT**TCC**GCGGCCGCTCGAGATCAGCCTCGA (forward) and 5’- TCGAGGCTGATCTCGAGCGGCCGC**GGA**ACGTGGCTTCTTCTGCCAAAG (reverse), creating a 13 aa long linker between the C-terminus of the PDGFR to the N-terminus of the FP, resulting in SNAP_mem_ (**Fig. 1a**).

#### SNAP-mCherry and variations in linker length

To examine how the length of the linker followed the stop codon, inserted between the two proteins, affects the expression of ORF2, td-tomato-C1 (SNAP*tdTom clone) was replaced by mCherry (Plasmid #238287 Addgene) using *XhoI*, and resulted in the ‘Original linker’ clone (**Suppl. 5a**). PCR amplification was done using the same primers as described above for td-tomatoC1 and e-GFP.

Next step was shortening the linker to a 3 aa long via a point mutation PCR using the primers; 5′-AAGAAGCCACGTTAG<u>GCGGCCGCG</u>ATGGTGTCTAAGGGTGAGGAA (forward) and 5′-TTCCTCACCCTTAGACACCAT<u>CGCGGCCGC</u>CTAACGTGGCTTCTT (reverse), resulted in the ‘Short linker’ clone (**Suppl. 5a**). For removing the linker and farther, removing the methionine, ‘Short linker’ clone served as the template for a reverse PCR performed with the following blunt primers; forward 5′-[P]**ATG**GTGTCTAAGGGTGAGGAA for the linker removal, and forward 5′- [P]GTGTCTAAGGGTGAGGAAGTA for the methionine deletion. For both PCRs, the same revers primer 5′-[P]CTAACGTGGCTTCTTCTGCCA was used. The linker deletion produced two different clones, a frame shifts mutant clone by the addition of a G nucleotide between the stop codon and the first Methionine named ‘frame shift’ and the wanted one ‘no linker’ (see **Suppl. 5a**). The methionine deletion resulted in the ‘no M’ clone (see **Suppl. 5a**). **KV_4.2_*GFP—**The eGFP clone (kindly provided by Prof. Aronheim A, Technion) was digested using *NotI/XbaI* and inserted following the TAA stop codon of the pCDNA kV4.2 plasmid (kindly provided by Alomone Labs™), resulting in KV4.2*GFP (**Suppl. 4a**). **GEFRI_1_—** SNAP*tdTom (**Fig. 1b)** served as backbone on which the first modification for preparing the GEFRI_1_, was done; replacing SNAP with GCaMP7s (kindly provided by Janelia Research Campus) via PCR amplification using the primers; 5′-TATATA**GAATTC**ATGGGTTCTCATCATCATCATCATCATGGTATGGC (forward) and 5′-TATATAAG**CGGCCG**CTCACTTCGCTGTCATCATTTGTACAAACTCTTC (revers). Cloning was done by *EcoRI/NotI* restriction and resulted with GCaMP*tdTom clone (**Suppl. Fig. 3a**). Second modification was done by cloning CFP (kindly provided by Prof. Dascal, Tel Aviv University, as reported^119^) as the first protein to be expressed, through ECoRI restriction (single digest enzyme reaction), which resulted in GEFRI_1_ (**Suppl. 6a, #1**) and flipped CFP (**Suppl. 8a**). PCR amplification was done using the following primers: 5′- TATACCG**GAATTC**ATGGTGAGCAAGGGCGAGGA (forward) and 5′- TATACCG**GAATTC**TCACTTGTACAGCTCGTCCATGC (reverse).

#### GEFRI_3_

To improve the expression of the two dORFs, we designed a Kozak sequence insertion (CCACC) prior to each ORF, following the stop codon, resulted in GEFRI_3_ (**Suppl. 6a, #3**). The CFP-GCaMP7s-tdTom were inserted into pAAV-CAG-eYFP (Plasmide #104055 Addgene), by replacing the YFP with the restriction enzymes XbaI (5’) and ECoRI (3’). The use of XbaI removed 114 bp from the 5`UTR of the plasmid. The clone was prepared by BioMatik company.

#### GEFRI_2_

For inserting back, the 114 bp to the 5’ UTR, CFP-GCaMP7s-tdTom - insert from GEFRI_3_ was cloned by BamHI/HindIII restriction into a pAAV CAG vector (Plasmid #100840 Addgene) by PCR amplification using the following primers: 5’- TATATACGCGGATCCTGTGACCGGCGGCTCTAGA (forward) and 5’- TATATATCGATAAGCTTGATATCGAATTC (reverse).

#### GEFRI_2_ construct preparation for stable cell line production

The backbone pSBbi-Pur (Plasmid #60523 Addgene) was amplified for inserting the restriction sites *HindIII* and *XBaI* using the following primers: 5’-TATATA**TCTAGA**AGGCCAGCTTGGGGTAGTTT (forward) and 5’-TATATA**AAGCTT**GCTCGCTGATCAGCCTCGA (reverse), respectively. GEFRI_2_ insert (CFP-GCaMP7s-tdTom) was cut out using *HindIII* / *XBaI* and inserted into the amplified plasmid, resulting in SB-GEFRI_2_. We co-transfected the SB-GEFRI_2_ (1.5 μg) with 0.5 μg of SB100- transposase (Plasmid #34879 addgene) to produced stable GEFRI_2_ HEK cells. The cells were supplemented with 5 μg/ml puromycin, and after 48hr seeded in 96 well plate for single-cell suspension, resulting in a single clone (**Suppl. 8**).

#### Split GFP varinats— Split sfGFP_1_

Split GFP was prepared by introducing a single stop codon in between GFP_1-10_ and GFP_11_ sequence fragments, using the well-known and useable split supper folder GFP (sfGFP). For this purpose, pCDNA 3.1-GFP_1-10_ (Plasmid #70219 Addgene) was used as the backbone to which a stop codon and a partial Kozak sequence together with the 11^th^ part of GFP were added by reverse PCR using the blunt primers; 5′- [P]GCTGGGATTACACTCGGCATGGACGAGCTGTACAAGTAAGGAACAGGTGGCGGC GGAA (forward) and 5′-[P]AGCATTTACATACTCATGAAGGACCATGTGGTCACGTCATTTTTCATTTGGATCTTTGC (reverse). However, those primers produced a mutant variant with an extra base at the fused spot, which destroyed the 11^th^ fragment. Therefore, we designed point mutation primers 5′-CCTTCATGAGTATGTAAATGCTGCTGGGATTACACTCGGCATGG and 5′-CCATGCCGAGTGTAATCCCAGCAGCATTTACATACTCATGAAGG to delete the extra base via PCR, resulted with **GFP_1*shift_** (**Suppl. 9b**). Subsequently, to improve the expression of the 11^th^ fragment, we added methionine prior to the 11^th^ part, by point-mutation PCR using the primers; 5’-CCAAATGAAAAATGACCATGCGTGACCACATGGTCC (forward) and 5’ – GGACCATGTGGTCACGCATGGTCATTTTTCATTTGG (revers), resulted in **GFP_1*M_** (**Suppl. 9b**).

#### Split-sfGFP_2_ variants

First, we introduced a stop codon with or without methionine between the 6^th^ to the 7^th^ strains of the sfGFP. We used sfGFP-N1 (Plasmid #54737 Addgene) as the backbone for inserting the stop codon with or without methionine following the N145 residue, via reverse PCR. We used the following blunt primers; 5′-[P] TTCAACAGCCACAACGTCTATATCACCGC (forward) or 5′-[P] ATGTTCAACAGCCACAACGTCTATATCACCGC (forward) with the 5′-[P] TCAGTTGTACTCCAGCTTGTGCCCCAGGAT (reverse), resulting in sfGFP_2*_ and sfGFP_2*M_, respectively (**Fig. 3a**). Then, we deleted 4 aa, following the N145 residue, between the 6^th^ to the 7^th^ strains of the sfGFP. We used sfGFP-N1 (Plasmid #54737 Addgene) as the backbone for inserting the stop codon and deleted the following FNSH residues via reverse PCR, using the blunt primers; 5′-[P] CACCAACGTCTATATCACCGCCGAC (forward) and 5′-[P] GCTCAGTTGTACTCCAGCTTGTGCCC (reverse), yielded sfGFP_2Del_. Next, methionine was inserted prior the 7^th^ strain by revers PCR using the blunt primers; 5′-[P] GAACGTCTATATCACCGCCGAC (forward) and 5′-[P] ATGGTGGCTCAGTTGTACTCCAG (reverse), resulted in sfGFP_2DelM_ (**Fig. 3a**). Finally, to separate sfGFP_2*M_ and sfGFP_2DelM_ into two plasmids, we used revers PCR using the primers; 5′-[P] GCGGCCGCGACTCTAGATCATAATCAGCCA (forward, sfGFP_1-6_), and 5′- [P]TCAGTTGTACTCCAGCTTGTGCCCCAGGAT (reverse, sfGFP_1-6_), to produce sfGFP_1-6_ (**Fig. 3c**), and 5′-[P] ATGTTCAACAGCCACAACGTCTATATCACCGC (forward, sfGFP_7-11_) or 5′-[P]AACGTCTATATCACCGCCGACAAGCAGAAG (forward, sfGFP_7-11Del_) with 5′-[P] CATGGTGGCGACCGGTGGATCCCGGGCCCG (reveres, sfGFP_7-11_), to generated sfGFP_7-11_ and sfGFP_7-11Del_, respectively (**Fig. 3c**).

#### Split-PA-sfGFP

PA sfGFP-N1 (Plasmid #54736 Addgene) was used as the backbone to separate it the same way as mentioned above, following the N145 residue via reverse PCR, using the blunt primers; forward_spGFP_1-6_, reverse_spGFP_1-6_, forward_spGFP_7-11_, or forward_spGFP_7- 11Del_, with the reveres_spGFP_7-11_, that used for clones sfGFP_1-6_, sfGFP_7-11_ and sfGFP_7-11Del_, resulting in the following clones; PAsfGFP_1-6_, PAsfGFP_7-11_ and PAsfGFP_7-11Del_, (**Fig. 3f**).

#### Split-GCaMP

GCaMP6s (addgene # 100843) was used as a template to engineer the GCaMP6s_1*-3.2*_.

#### GCaMP6s_1*_

Stop codon was inserted at position 27 (after L25E26 linker) via PCR for engineering GCaMP6s_1*_ (**Suppl. 12a**) using the primers 5′- GGTCGGCTGAGCTCACTCGAGTAGAACGTCTATATCAAGGCCGAC (forward) and 5′- GTCGGCCTTGATATAGACGTTCTACTCGAGTGAGCTCAGCCGACC (reverse).

#### GCaMP6s_2*_ variants

To create super folder GCaMP6s and optimal super folder GCaMP6f we based on the PA-GCaMP6s/f^98^. First to reveres engineer the photo-activity we exchange F221L and S222T using a point-mutation PCR using the following primers: 5’- CCCACCCTCGTGACCACC**C**T**G**A**C**CTACGGCGTGCAGTGCTTCAGC (forward) and 5’ – GCTGAAGCACTACACGCCGTAG**G**T**C**A**G**GGTGGTCACGAGGGTGGG (reverse), and the D115V, with the primers; 5’- CCCGACAACCACTACCTGAGCG**TG**CAGTCCGTGCTTTCGAAAGAC (forward) and 5’ – GTCTTTCGAAAGCACGGACTG**CA**CGCTCAGGTAGTGGTTGTCGGG (reverse). Finally, to engineer the split GCaMP6s_2*_, sfGCaMP6s_2*_ and osfGCaMP6f_2*_ (**Suppl. 12a** and **Suppl. 13a,** respectively) we introduced a stop codon with a partial Kozak sequence [C] prior the second part of the cpGFP, replacing the serine in the linker with stop codon, through revers PCR using the primers; 5’- ACATGGTGAGCAAGGGCGGTACCGGAGGGTG (forward) and 5’ – CACCCTCCGGTACCGCCCTTGTACAGCTCGTCC (reverse).

#### Split PA-GCaMP6s

The sfGCaMP6s_2*_ used as a template to engineer split PA-sfGCaMP6s_2_ (**Suppl. 14a**). We exchanged the V115H, L221F and T222S first by amplification of three fragments using three sets of primers: hSyn promoter_F 5`- CGCACCACGCGAGGCGCGAGATAGG (forward) and 5`- GGGGTCTTTCGAAAGTTTGGACTG**GTG**GCTCAGGTAGTGGTTGTCGGGC (reverse) to amplify the fragment between the human synapsin promoter to the mutagenesis V115H site, the 5`-GCCCGACAACCACTACCTGAGC**CAC**CAGTCCAAACTTTCGAAAGACCCC (forward) and 5`-GGCTGAAGCACTGCACGCCGTAG**C**T**G**A**A**GGTGGTCACGAGGGTGGGCCAG (reverse) used to amplified the fragment between the two sites of mutagenesis, and the point mutation primer 5`- CTGGCCCACCCTCGTGACCACC**T**T**C**A**G**CTACGGCGTGCAGTGCTTCAGCC (forward) and 5`- TCACTTCGCTGTCATCATTTGT(reverse) to amplified the part between the mutagenesis sites L221F and T222S to the end of the CAM. Next, the three fragments were assembled by Gibson assembly kit (NEB E2611), followed by PCR amplification using the two distal primers: hSyn promoter_F 5`-CGCACCACGCGAGGCGCGAGATAGG (forward) and the CAM_R 5`- TCACTTCGCTGTCATCATTTGT (reverse) to amplified only the assemble fragment, starting from hSyn promoter and ending at the CAM. Finally, the amplified product, and the vector, pAAV-CAG-eYFP (Plasmide #104055 Addgene), were digest with NheI and PstI, ligated, and transform into bacteria (for more information see^120^).

#### Split-GCaMP6s_2*_ variants

Finally, to separate GCaMP6s_2*_, sfGCaMP6s_2*_ and osfGCaMP6f_2*_ into two plasmids, we used revers PCR using the primers; 5′- GCCACCATGGTGAGCAAGGGCGAGG (forward, GCaMP_1-6_) with 5′- CTTGCTCACCATGGTGGCGAGATCTGAGT (reverse, GCaMP_1-6_), or 5′- ACCGGAGGGTGAACGGCCGCGACTCTA (forward, GCaMP_7-11_) and 5′- CGGCCGTTCACCCTCCGGTACCGCC (reveres, GCaMP_7-11_), and assemble the fragments using Gibson assembly (NEB E2611) to generated GCaMP_1-6_ or GCaMP_7-11_ parts (**Suppl. 13a**).

#### GCaMP6s_3*_ variants

A stop codon was inserted prior the L268-P269 linker residue using the following primers: 5′- CACAAGCTGGAGTACAACTGACTGCCGGACCAACTGACTGAAGAG (forward) and CTCTTCAGTCAGTTGGTCCGGCAGTCAGTTGTACTCCAGCTTGTG (reverse), resulting in GCaMP6s_3*_ (**Suppl. 12a**). Next, L268 in GCaMP6s was replaced with a stop codon using the following primers: 5′- GGGCACAAGCTGGAGTACAACTGACCGGACCAACTGACTGAAGAG (forward) and 5′- CTCTTCAGTCAGTTGGTCCGGTCAGTTGTACTCCAGCTTGTGCCC (reverse), resulting in GCaMP6s_3.1*_ (**Suppl. 12a**). Similarly, for producing GCaMP6s_3.2*_ (**Suppl. 12a**), P269 was replaced with a stop codon using the primers as follows: 5′- GGGCACAAGCTGGAGTACAACCTGTGAGACCAACTGACTGAAGAG (forward) and 5′- CTCTTCAGTCAGTTGGTCTCACAGGTTGTACTCCAGCTTGTGCCC (reverse).

#### Split GCaMP6s_3*_ into M13-cpGFP and MLP-calmodulin

GCaMP6s_3**\***_ was used as a template to separately amplify M13-cpGFP or calmodulin fragments. First, M13-cpGFP (**Suppl. 12** and **Suppl. 18**) was amplified via revers PCR using the primers: 5′-[P] TCAGTTGTACTCCAGCTTGTGCCCCAGGAT (forward) and 5′-[P] TCAGTTGTACTCCAGCTTGTGCCCCAGGAT (revers). After which, MLP-CaM fragment (**Suppl. 12**) was prepared similarly using the following primers: 5′-[P] TTCCGGGTTGTACTCCAGCTTGTGCCC (forward) and 5′-[P] TTCCGGATGCTGCCGGACCAACTGACTGAA (reverse).

#### Split jR-GECO_1a_

Split jR-GECO_1a*120_ (**Suppl. 15**) was generated on the base of jR**-**GECO_1a_ (kindly provided by Janelia Research Campus) construct via PCR amplification using the SDM using the primers as follows: 5′- GGAGGTACAGGCGGGAGTTGACTGGTGAGCAAGGGCGAG (forward) and the 5`-CTCGCCCTTGCTCACCAGTCAACTCCCGCCTGTACCTCC (revers), for inserting TGA stop codon following the linker of cp-mApple (S120). Then, Split jR-GECO_1a*120_ was separated via revers PCR with the primers as follows: 5′-[P] TAGGCGGCCGCGACTCTAG (forward) and reverse 5′-[P] ACTCCCGCCTGTACCTCCC, for generating Split jR-GECO_1a_-part I, and 5′-[P] CTGGTGAGCAAGGGCGAGGA (forward) and the 5′-[P] CATGGTGGCGAGATCTGAGTC (revers), resulting in Split jR-GECO_1a_-part II (**Suppl. 15**).

### Labeling Mito-ER contact sites by MEGIC variants

#### MEGIC_1_

First the calmodulin fragment was fused to CB5 (ER transmembrane domains), to do that, xbaI restriction site sequence (TCTAGA) was inserted after the calmodulin in clone GCaMP6s_3*_ using blunt primers: 5′-[P]AGATGAAAGCTTATCGATAATCAACCT (forward) and 5′-[P]AGACTTCGCTGTCATCATTTGTACAA (reverse). In parallel, CB5 (including the linker prior it; SAGGSAGGSAGGSAGGSAGGPRAQASNSSAGG) was amplified with xbaI site from the backbone plasmid (plasmid #120915 addgene) using the primers; 5’- TATATAGCTCTAGAAGTGCTGGTGGTAGTGCTGG (forward) and 5’-TATATAGCTCTAGAGTCCTCGGCCATGTACAGGC (reverse), resulted in the subclone GCaMP6s_3*_-CB5. Next, SphI restriction enzyme site sequence (GCATGC) was inserted to GCaMP6s_3*_-CB5) using blunt the primers: 5′-[P] TGCATGGTCGACTCATCACGTCG (forward) and 5′-[P] TGCGGTGGCGAGATCCTTATCGT (reverse). Likewise, TOM (including the linker GSGASGSAGGGV) was amplified from (plasmid #120914 addgene) with SphI enzyme restriction site using the following primers: 5’- TATATA**GCATGC**ATGGTGGGTCGGAACAGCGC (forward) and 5’-TATATA**GCATGC**CACTCCACCACCAGCACTTC (reverse) and fused before M13-cpGFP, resulting with Tom-GCaMP6s_3*_-CB5. Then, the Tom-GCaMP6s_3*_-CB5 was amplified with AgeI using blunt primers; 5′-[P] GGTTGAAAGCTTATCGATAATCAACCT (forward) and 5′-[P] GGTTCTAGAGTCCTCGGCCATGTACAG (reverse). In parallel, mRuby3 was amplified from its backbone (plasmid #112005 addgene) inserting *AgeI* restriction site using the primers: 5′- TATATA**ACCGGT**ATGGTGTCTAAGGGCGAAGAG (forward) and 5′-TATATA**ACCGGT**TCACTTGTACAGCTCGTCCATGC (reverse) and cloned to Tom-GCaMP6s_3*_-CB5, resulting in Tom-GCaMP6s_3*_-CB5-mRuby3.

#### Tom-M13cpGFP

Tom-M13cpGFP (clone #50, suppl. Table 1) was amplified from Tom-sGCaMP6s_3*_-CB5 backbone with the same primers and PCR conditions used to amplify M13cpGFP (see above; **Suppl. 18**).

#### CAM-CB5-mRuby3

To produced CAM-CB5-mRuby3 (**Suppl. 18**), was amplified from Tom-GCaMP6s_3*_-CB5-mRuby3 clone using the following blunt primers: 5′-[P] GCCACCATGCTGCCGGACCA (forward) and 5′-[P] GGTGGCGAGATCCTTATCGTCATC (revers).

### Labeling Mito-ER contact sites using split-sfGFP

To produced TOM20-GFP_1-6_ we amplified Tom-M13cpGFP via revers PCR, using the primers: 5′- GCACAAGCTGGAGTACAACTG (forward), and TCCTCGCCCTTGCTCACCATGCATGCCACTCCACCA (revers). In parallel, sfGFP_1-6_ was amplified from the previous sfGFP_1-6_ (**Fig. 3c**) using the primers: 5′- TGGTGGAGTGGCATGCATGGTGAGCAAGGGCGAGGA (forward), and 5′-CAGTTGTACTCCAGCTTGTGC (revers). Both fragments were digested using Gibson assembly (NEB E2611), resulting in TOM20-GFP_1-6_ (**Suppl. 16**). Similarly, we amplified the CB5-mRuby3 from the previous CAM-CB5-mRuby3 clone via revers PCR, using the primers: 5′- GACGAGCTGTACAAGTCTAGAAGTGCTGGTGGTA (forward), and 5′- ACCAGCACTTCTAGACTTGTACAGCTCGTCCATGC (revers). In parallel, sfGFP_7-11_ was amplified from the previous sfGFP_7-11_ (**Fig. 3c**) using the following primers: 5′- GGATCCACCCGCCACCATGAACGTCTATATCA (forward), and 5′-ACTTGTACAGCTCGTCCATGC (revers). Both fragments were digested using Gibson assembly (NEB E2611), yielding GFP_7-11_-CB5-mRuby3 (**Suppl. 16**).

#### MEGIC_2_

We designed primers which were compatible for GCaMP6s_2*_, sfGCaMP6s_2*_ and osfGCaMP6f_2*_. The first parts (wt-sf-and osf-TOM-M13-GFP_7-11_) were produced by amplifying the GFP_7-11_ using the following primers: 5’- TCGGCTGAGCTCACTCGAG (forward), and 5′- CCC**AAGCTT**TCACTTGTACAGCTCGTCCATG (revers) inserting *HindIII* restriction site to the 3’-end. Next, the amplified inserts and the TOM-M13-cpGFP plasmid were digested by *XhoI/HindIII* and ligated for yielding parts I of wt-sf-and osf-MEGIC_2_ (**Suppl. 18**). To generate the parts II of MEGIC_2_ (wt-sf-and osf-GFP_1-6_-CaM-CB-mRuby3; **Suppl. 18**), we used Gibson assembly (NEB E2611) to produce all the clones by amplified the inserts (wt-sf-and osf-GFP_1-6_) using the following primers: 5′-GGATCCACCCGCCACCATGGTGAGCAAGGGCGAGGAGC (forward), and 5′-TTCAGTCAGTTGGTCCGGCAG (revers), and the backbone CaM-CB-mRuby3 (**Suppl. 18**) using the primers as follows: 5’- GCTCCTCGCCCTTGCTCACCATGGTGGCGGGTGGATCC (forward), and 5′- GGCACAAGCTGGAGTACAACCTGCCGGACCAACTGACTGA (revers).

### Split-sfGCaMP6s_2_ clone for AAV production

We subcloned split-sfGCaMP6s_2_ (from pGP-GCaMP6s-^27^) to another AAV compatible plasmid (Addgene, Plasmid#162374). We removed the M13-sfGFP_1-6_ fragment by digesting it from clone TOM-M13-cpGFP (**Suppl. 17**) using *BamHI*. For preparing part II, we amplified the sfGFP_7-11_- CAM using the following primers: 5′-TATATACGC**GGATCC**CAGATCTCGCCACCATGGTG (forward), and 5′-TATATACCC**AAGCTT**CTCACTTCGCTGTCATCATTTG (revers), digested with *BamHI/HindIII* and cloned to the *BamHI/HindIII* digested pGP-AAV-syn-jGCaMP8s (Plasmid #162374 Addgene).

### Zebrafish constructs

Split-sfGCaMP_2_ parts; M13-sfGFP_7-11_ and sfGFP_1-6_-CAM were amplified each from their backbone using the following primers 5`- TATATA**ACCGGT**GCCACCATGGGTTCTCATCAT (forward) and 5`- TATATA**ACCGGT**AGAGTCGCGGCCGTTCACC (reverse) or 5`- TATATA**ACCGGT**ATGGTGAGCAAGGGCGAGG (forward) and 5`-TATA**ACCGGT**CTCACTTCGCTGTCATCATTTG (reverse). Following amplification, fragments were digested by *AgeI*, as this was the restriction site compatible in the ZF vector – HuC (kindly provided by Prof. Kawashima T). Likewise, Tom-M13-sfGFP_7-11_ were amplified using the primers:5`- TATATA**ACCGGT**ATGGTGGGTCGGAACAGCG (forward) and 5`- TATATA**ACCGGT**TCACTTGTACAGCTcGTCCAT (reverse), digested with *AgeI* and inserted to the HuC vector. The sfGFP_1-6_-CAM-CB5-mRuby3 were inserted using Gibson assembly (NEB E2611). The insert was amplified with the following primers: 5`- TGCAGATAATTGCCACCATGGTGAGCAAGG (forward) and 5`-CAGGATCCACCGGTTCACTTGTACAGCTC (reverse), and the HuC vector by 5`- AAGTGAACCGGTGGATCCTGGCCG (forward) and 5`- ACCATGGTGGCAATTATCTGCAGGTGGAAAATATA (reverse).

### Cells cultures

#### Heterologous cell lines

HEK293T and HeLa cells, were grown in 10 cm^2^ cell culture plate (corning) at 37°C with 5% CO2 in Dulbeccòs modified Eaglès medium (DMEM-F12) supplemented with 10% fetal bovine serum (FBS) and 1% of L-glutamine (full media). <u>Splitting</u> <u>HEK293T and HeLa cells</u>-The cells were washed with 2 ml of PBS X1 twice after 4 days of incubation to remove FBS protein. For suspension of the HeLa cells, 1 ml of trypsin was added while tapping and incubated for 2 minutes to float the cells. After incubation, 10 ml of medium was added gradually to counteract the trypsin while pipetting slowly. The cells in the medium were split into new 10 cm2 dishes.<u>DNA transfection</u> HEKand HeLa cells were grown to ∼50% confluence on poly-D-lysine (PDL) coated 12 mm coverslips (Bar-Naor, Israel) and incubated in full media at 37°C with 5% CO2 in 24 well plates overnight prior transfection. Cells were transfected with 1 µg of DNA diluted in 45 µl dMEM and then mixed with 3 µl ViaFect™ (Promega, USA) transfection reagent added and incubate for 20 minutes. The diluted DNA was incubated with the HEK or HeLa cells for 24-48 hours prior imaging.

#### Hippocampal cells culture

Primary hippocampal neurons were extracted from Spague-Dawley rat pups (P0-1) as previously described^121^. Briefly, hippocampal neurons were grown on PDL (1mg/ml diluted in DDW) coated 12 mm glass coverslips and transfected in between day 7-10 from extraction. Transfection was done with 1 µg DNA using the calcium-phosphate transfection method (Kingston et al. 2001) and imaged at 48-96 hours post transfection. All animal procedures were approved by the Technion’s Institutional Animal Care and Use Committee (permit no. IL130- 09-17).

### SNAP-tags for decorating cells by orthogonal labeling

SNAP-Surface® Alexa Fluor® 488 (NEB) were dissolve in DMSO to yield a labeling stock solution of 1 mM tags’ substrate. Mixed by vertexing for 10 minutes until all the tags’ substrate dissolved completely. The 1 mM stock solutions were diluted 1:4 (v/v) in fresh DMSO to yield a 250 µM stock for labeling proteins in vitro. The stock labeling was diluted 1:250 in fresh full media and incubated with the cells protected from light, for 30 minutes at 37°C with 5% CO2. Cells were washed x3 using complete media to remove nonspecific binding.

### Transgenesis of Zebrafish larvae and imaging

At day of fertilization, embryos were injected with a mixture solution containing; 10 µL of 0.4 M KCl, 2 µL of 250 ng/µL transposase mRNA, 2 µL of 250 ng/µL plasmid DNA (containing the above two parts of split ZF-sfGCaMP6s_2_ or separated ZF-MEGIC_2_ plasmids), and 6 µL of water to reach a final volume of 20 µL. Injected embryos were raised to day 5-7, reared in E3. On days 5-7 larvae were embedded in 1.4% low-melting agarose in E3 and soaked in E3 for an imaging session using confocal LSM 900 (Zeiss, Germany). E3 medium working concentration: 5mM NaCl, 0.17mM KCl, 0.33mM CaCl_2_, 0.33mM MgCl_2_.

### Fluorescence Imaging

Imaging was performed on a Zeiss Laser Scanning Confocal Microscope equipped with a spectral detector and Airysan2 (LSM-880 and LSM900, Zeiss, Germany, respectively). Cells were imaged in a standard clear imaging medium containing (in mM): 138 NaCl, 1.5 KCl, 1.2 MgCl2, 2.5 CaCl2, 10 D-glucose, 10 HEPES, pH 7.4., with a water Plan-Apochromat objective lens (20x/1.0 DIC D=0.17 (UV) VIS-IR M27 75 mm, ZEISS). Excitation of the red fluorescent proteins; td-Tomato, mcherry, mRuby and jRGECO was performed by 651 nm diode laser, the GFP and GCaMP variants constructs by argon ion laser (488 nm) while the CFP by 405 nm diode laser. Emission spectra were collected using a spectrally resolved 32-pixel GaAsp detector array.

<u>For Ca^2+^-imaging</u> using GCaMP variants, final concentrations of ionomycin and Ca^2+^ were 10nm and 20 mM, respectively, were added to the media while imaging, to detect the changes of fluorescents in real time.

<u>Photoactivation</u>—Image acquisition and photoactivation (i.e., bleach mode; Zen software, Zeiss) performed intermittently, acquired between pulses of 405 nm illuminations. Time resolved fluorescence plots thereby present data from images acquired before and after the 405 nm photoactivation bouts.

### Fluorescence activated cell sorter (FACS) experiments

Expression of the different fluorescent proteins (ORFs) of the stable GEFRI_2_ line were measured by High Throughput LSR Fortessa (BD Biosciences) fluorescence-activated cell sorter (FACS). V-450/30, B-530/30 and YG-586/15 (band pass) filters were used for the CFP, GCaMP7s and td-Tom, respectively.

### Electrophysiology

Patch-clamp recordings were performed 24-48 hours after transfection of HEK cells or at 16-23 DIV (days *in vitro*) for dissociated hippocampal neurons, as previously described^122^. Briefly, patch-clamp recordings were performed with an Axon MultiClamp 700B amplifier and Axon Digidata 1440A acquisition system, mounted on an OLYMPUS BX51WI microscope (equipped with 5× and 40× objectives) for split-GCaMP6s and Kv4.2 constructs. Patch-clamp recordings of neurons expressing split-MEGICs were performed using the loose patch method^27^, with an Axon AXOPATCH 200B amplifier system and Axon Digidata 1440A mounted on a LSM900 confocal microscope. Illumination was performed using a TH4-200 microscope light power supply (OLYMPUS) and X-Cite® 110LED (EXCELITAS Technologies) with 475 LEDs for split GCaMP3 and Kv4.2 constructs and LSM900 lasers for split-MEGIC constructs. Imaging during patch-clamp recordings was performed using µManager^123,124^ with a Pixelink® camera sensor for split-GCaMP6s. All cells were recorded with a low-pass filter at 10 kHz. Pipette resistance ranged between 5 to 11 MΩ for whole cell recordings (split GCaMP3 and Kv4.2 constructs) and 3 to 6 MΩ for cell attached recordings (MEGIC constructs) and were filled with an internal solution containing (in mM): 135 K-Gluconate, 10 NaCl, 10 HEPES, 2 MgCl2(H2O)6, 1 EGTA, 2 Mg-ATP, pH 7.3 with KOH.

<u>HEK293t cells</u> were clamped at -70 mV, and full IV curves were typically obtained by a series of 200-400 ms long voltage steps, ranging from -100 or -50 mV to +60 mV, in 10 mV increments with each step lasting 200 ms. For HEK cell recordings, the extracellular recoding solution contained (in mM): 135 NaCl, 4 KCl, 10 HEPES, 10 D-glucose, 1.8 CaCl_2_(H_2_O)_2_, 1 MgCl_2_(H2O)_6_, pH 7.4 with NaOH. IV-curves for all Kv4.2 channel recordings was calculated as follows: the peak current after the initial stimuli (5-30 ms) was subtracted by the leak current towards the end of the stimuli (Kv4.2 is a type-A voltage-gated channel, i.e., fast deactivating, therefore shows no residual current past 200 ms). <u>Neurons</u> were clamped at -70 mV for voltage clamped or left at I=0 for current clmap recordings, recorded under the gap-free mode. Neuronal extracellular recording solution consisted of (in mM): 138 NaCl, 1.5 KCl, 5 HEPES, 10 D-glucose, 2.5 CaCl_2_(H_2_O)_2_, 2 glycine, pH 7.4 with NaOH.

#### Electrophysiology Analysis

All analyses described were performed using the Clampfit software (Molecular Devices) and plotted (and tested statistically) using GraphPad Prism 8. Number of Action potentials (#Aps) were manually counted and correlated with Δ*F/F* recordings. The #APs vs. Δ*F/F* were fitted using a standard exponential function (GraphPad Prism 8).

### Western Blot Analysis

HEK293t cells were grown to ∼70% confluency on 10 cm^2^ plates (Corning) at 37°C with 5% CO_2_ in full media (as mentioned above). Cells were transfected with 12 µg of DNA diluted in PEI solution (36 µl of PEI, 1mg/ml, in 0.5 ml PBS) and incubated for 48 hours at 37^0^C with 5% CO_2_ prior harvesting. Protein extraction and blotting was performed 24 - 48 hours post transfection. HEK cells were washed three times with ice-cold PBSx1 and lysed using Maltoside lysis buffer [1% *N*-dodecyl maltoside (cat no. 69227-93-6 – Sigma) dissolved in PBSx1 with freshly added 0.1 mg/ml PMSF and protease inhibitors], incubated on ice for 1 hour with vertexing each 15 min followed by centrifugation at 16,000 X g for 30 min. Proteins were then separated by 10% SDS-PAGE and transferred onto a nitrocellulose membrane. Membranes were incubated in 3% non-fat dry milk in PBSx1 for 1 hour at RT, and subsequently probed with one of the primary antibodies for 16 hours at 4^0^C. Primary antibodies used; Anti tdTom (ChromoTek™ #5f8 mAb) and anti myc (purified and concentrated from 9E10 hybridoma cell line generously provided by Prof. Ami Aronheim – Technion). Primary antibodies were detected using the HRP-conjugated Donkey-anti-rat or anti-mouse (Jackson ImmunoResearch Laboratories, USA) secondary antibodies. The detection was done by *Enhanced chemiluminescence* (*ECL*).

### AAV production

Transfection and virus collection-HEK293-T were seeded on six 10 cm^2^ plate dishes in DMEM-F12 supplemented with 10% FBS and 1% glutamine. 24h after plating, transfection was performed using PEI (Polyethyleneimine 1mg/ml in PBSx1) and three plasmids: pAAV Helper which encodes adenoviral proteins necessary for replication, pAAV Rep-Cap which encodes the viral replication and capsid proteins, in a 3:2:5 molar ratio, respectively (∼27µg DNA/plate). 6-8 h post-transfection old media was changed to a new DMEM-F12 media supplemented only with 1% glutamine. 48- and – 72h post transfection media containing pAAV viruses was collected and filtrated through a 0.45µm filter. Viruses were also collected from lysed transfected-HEK cells in gradient buffer (described in Materials), undergo 3 rounds of freeze/thaw cycles in liquid nitrogen and a 37^0^C water bath, followed by triturating through an 18G needle for 8-10 times. Virus cells lysate was centrifuged at 3,724xg and 4^0^C for 10 min, and filtered through a 0.45µm filter. Virus concentration-For first round of concentration, gradient buffer x10 (described in Materials) was added to viral sup media followed by centrifugation using Centricon Plus-70 (UFC703008 - Millipore) for ending-up with ∼4-6 ml of viral media (at this stage virus could be kept at 4^0^C for over-night or at -80^0^C for longer storage). For destroying left plasmid-DNA from transfection, viral media was treated with Benzonase (E1014-25KU 250Units/ul – Sigma, at 1µl/5ml viral media, for 1h at 37^0^C water bath), followed by second round of concentration by loading on an Iodixanl-density gradient media (OptiPrep 60% w/v D1556 – Sigma, below), and centrifuge in a Beckman WX Ultra90 centrifuge at 41,000rpm and 18^0^C for 2.5h. After centrifugation, virus was collected from the 40% Iodixanol layer, washed (at 3,724xg at 4^0^C for 3-5 min) (x3) with PBSx1 using Amicon Ultra-15ml (UFC910024 - Millipore), followed by concentration in storage buffer (described in Materials) using Amicon Ultra -0.5ml (UFC510024 - Millipore) and stored at -80^0^C. Virus titer was tested by qPCR using the forward; 5’-GCTGTTGGGCACTGACAAT-3’ and reverse; 5’-CCGAAGGGACGTAGCAGAAG-3’ -WPRE primers. Materials for gradient:

- Gradient buffer x10: 100mM Tris (pH 7.6), 1.5M NaCl, 100mM MgCl_2_, Filter sterilizes using 0.22 µM Vacuum filter.
- Storage buffer (for 1 litter): 50gr D-Sorbitol, 212mM NaCl - diluted in PBSx1.
- Iodixanol (OptiPrep 60% w/v D1556 - sigma) solutions for gradient preparation:

|  | 15% +1M NaCl | 25% | 40% | 58% |
| --- | --- | --- | --- | --- |
| Optiprep 60% (ml) | 40ml | 46.67 | 64 | 96.67 |
| x10 GB (ml) | 16 | 11.2 | 9.6 | 3.33 |
| 5M NaCl (ml) | 32 | 0 | 0 | 0 |
| ddH<sub>2</sub>O MilliQ water (ml) | 72 | 56 | 22.4 | 0 |
| Phenol Red (5mg/ml) | 0 | 280ul | 0 | 240ul |

Loading is done from lightest to heaviest in a 11.2ml tube (Ultra-Clear 344059 – Beckman).

### Infection GEFRI_2_ in vivo

1-2 days old Sprague Dawley rats (males or females from Envigo, Israel) were infected by injection of AAV (∼4µl of ∼ 1×10^12^ vg/ml) to their left hemisphere. After 1-2 months old rats undergo trans cardiac perfusion with PBSx1 (02-023-5A, Biological Industries) followed by neutral buffered 10% formalin (HT5011, Sigma-Aldrich). The whole brain was removed and immersed in 4% paraformaldehyde (PFA) for ∼16 hours at 4^0^C. For <u>cryosection preparation</u> brains were sequentially immersed in 15%, 20% and 30% sucrose in PBS, for 1 hour at room temperature with each of the first two sucrose solutions, and for ∼16 hours in the 30% solution at 4^0^C. Then brains were embedded in OCT (Optimal Cutting Temperature compound) and kept at -80^0^C for ∼16 hours prior to frozen sectioning on a microtome-cryostat. Tissue sections were cut at 16µm thickness and mounted on cover slips (supper-frost plus, Thermo) followed by confocal – imaging (**Fig. 2**).

### Surgical preparation

Surgical procedures were performed under isoflurane anesthesia (4% for induction and 1.5-2% during surgery). Mice aged 8-12 weeks were anesthetized and secured in a stereotaxic apparatus. A heat pad was used to maintain body temperature at 36-37 °C. The scalp was shaved and cleaned with iodine solution and ethanol. The skull surface was exposed after a subcutaneous injection of 2% Lidocaine. A small circular craniotomy (2.5-3 mm diameter) was performed for viral injections over the primary motor cortex forelimb area 0.6 mm anterior and 1.6 mm lateral to Bregma. 50 nL of the viral mixture (AAV1-TOM20-M13-cpGFPII; AAV1-cpGFPI-CaM-CB5-mRuby3= Split-MEGIC_2_) was injected using a hydraulic micromanipulator (M0-10 Narishige) at several sites within the craniotomy at 300 µm below dura. After the injections, an optical window constructed from a 2.5 mm and 3.5mm 170 µm thick circular glass disks glued together was placed over the exposed brain and superglued in place. A custom-made 3D printed headpost was affixed to the skull using dental cement.

Injections of ketoprophen (5 mg/kg) and buprenorphine (0.1 mg/kg) were administered subcutaneously for analgesia during surgery and for 2-days post operation. Mice recovered for at least one week following surgery with *ad libitum* food and water.

### Two-photon imaging and data analysis

Images were acquired using a two-photon microscope (Bruker 2P-Plus) as previously reported^125^.Briefly, we used a 2P-micrsocope equipped with a dual-beam Insight X3 laser (Spectra Physics), an 8 kHz bidirectional resonant galvo scanner and a Nikon 16X CFI Plan Fluorite objective (NA 0.8), controlled by the software package PrairieView 5.3. Fluorescence was split by a 565LP dichroic and filtered with 525/70 and 595/50 bandpass filters before collection on two GaAsP photomultiplier tubes (Hamamatsu H10770PB-40 and H11706-40, respectively) to image the red (mRuby3) and green (split-MEGIC_2_) fluorescent markers. The imaging region was ∼400 x 400 µm, 512×512 pixels at a frame rate of 30 Hz. Illumination was centered at a wavelength of 940 nm with 30-40 mW mean power at the objective. PMT gains were set to minimize saturated pixels during calcium transients. Images were acquired in segments of 60 sec followed by an inter-acquisition interval of 15 sec. The fluorescence data acquired by the two-photon microscope were first registered to correct for brain motion artifacts. Our registration method was based on Fienup, J. & Kowalczyk, using Fourier transform-based correlation between two successive images. The maximal value position in the correlation image specifies the relative shift between the two images; we designate them *u_t_* and *v_t_*. This method required a template specification and matching against an image stack. The template image *I_temp_*(*x*, *y*) was defined as the average of all images in the selected trial over time.

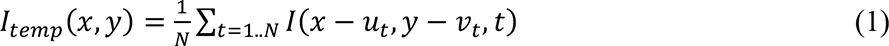

The set {*u_t_*, *ν_t_*}, *t* = 1. . *N* is an image shifted in the XY plane after alignment. We initially started with *u_t_=0, v_t_=0* and then updated their values according to the registration maxima. This procedure was repeated several times, when each time we computed the new template *I_temp_*(*x*, *y*) using previously computed {*u_t_*, *ν_t_*} for each image. Typically, this procedure converges after several iterations, in our case three iterations. To align the imaging data over many trials, we used a similar technique, utilizing the previously computed averaged templates for each trial. For each trial *k*, we performed a single trial registration using the template algorithm for three iterations. To align the image data over many trials, we treated the final templates 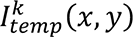 from each trial *k* as unaligned image data and repeated the same registration procedure to find offsets {*u_k_*, *ν_k_*} for each trial. These offsets, along with previously found offsets {*u_t_*, *ν_t_*}, account for the final image shift in the XY plane.

Regions of interest (ROIs) were detected manually using average fluorescence images and ΔF/F projection images, which highlighted active neurons. The pixels within each ROI were averaged for every frame. The ROI "mask" was used to detect the same neurons on multiple imaging sessions on different days.

*ΔF/F* was computed using the following formula:

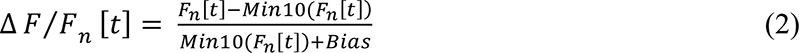

*Min10(Fn [t])* is a mean value of the lowest consecutive 10% values of the fluorescence signal *Fn [t]*. This minimal fluorescence value was calculated both per trial and across the entire experimental session, with no significant differences in the results with these two calculation variants. A small bias factor in the denominator prevented division by zero when the cell was completely silent.

### Experimental models

All animal procedures were performed in accordance with guidelines established by the NIH on the care and use of animals in research, as confirmed by the Technion Institutional Animal Care and Use Committee. Adult C57BL mice were used in this study. Animals were housed in a 12:12 hours reverse light:dark cycle.

### Data analysis

Statistical analysis change in fluorescence (Δ*F/F*) was calculated by (F_t_–F_0_)/F_0_, where Ft is the measured fluorescence (in arbitrary units, a.u.) at a given time t, and F_0_ is the initial baseline fluorescence, typically calculated from averaging the baseline. Δ*F/F* - 1 describes an increase by 100%, equivalent to two-fold increase in fluorescence. Owing to easily observable basal fluorescence, we did not need encounter near division-by-zero artifacts. For collecting emission spectra, we used small imaging intervals (8 nm, see above) which allowed us to plot the data as smooth (splined) curves rather than straight lines (SigmaPlot 11). <u>Statistical analysis</u>: Results are shown as mean ± SEM. Two group comparison were performed using non-parametric Mann-Whitney test and multiple group comparison were performed using 2-way ANOVA Tukey multiple comparison test (GraphPadPrism9). Correlations between two independent parameters were examined using Spearman correlation, by Kolmogorov-Smirnov comparison tests. Asterisks indicate statistically significant differences: *, P < 0.05; **, P < 0.01; ***, P < 0.001.

---

**Supplementary figure 1. Stop codon-mediated bicistronic expression of two independent proteins with diverging subcellular localizations—** Micrographs of HEK293 cells co-expressing SNAP-tdTom (**a, cartoon**) or SNAP*tdTom (**b, cartoon**) clones (tdTom, magenta) and an ER marker-SEC61 GFP (green). Introduction of a stop codon before the ORF of tdTom, completely separates the fluorescent protein from its parent protein (a membrane tethered protein, also appearing in ER) and therefore appears to fill entire cytoplasm. These are summarized in the fluorescence intensity profiles shown below micrographs (trajectories of the plot are depicted by dashed white lines in bottom-left micrographs). Note how the presence of a stop codon completely alters the colocalization patterns of magenta (tfTom) and green (ER) traces.

**Supplementary figure 2. Stop codon-mediated bicistronic expression in multiple cells lines—a.** Micrographs of HeLa cells expressing the SNAP-tdTom (left) and SNAP*tdTom (right) clones. tdTom fluorescence (magenta) appears intracellularly and colocalizes with the membrane labeling of SNAP-tethered (by BG-Alexa488, green) appearing in white. Top two HeLa cells shows strong colocalization at the membrane surface of lamellipodia. (right) Introduction of a stop codon separates between the two proteins, abolishing colocalization (note lack of white overlap). **b.** Representative micrographs of primary cultured neurons exhibiting bicistronic expression when transfected with SNAP*tdTom (magenta) and labeled by BG-Alexa488 (green).

**Supplementary figure 3. Stop codon-mediated bicistronic expression of varyiong ORFs, including GCaMP, in multiple cell lines— a, b.** Illustration (**a**) and micrographs of HEKs cells (**b**) expressing a bicistronic clone containing GCaMP7s as ORF1 (green) followed by stop codon (red, UGA) and tdTom (ORF2). (right) Summary of the fraction of cells in which both ORFs could be detected. **c, d.** Bicistronic expression of clone shown in (**a**) in glia (**c**) and neurons (**d**). (**d,** left) Representative traces showing Ca^2+^-activity reported by GCaMP7s. **e.** Cartoon (left) and micrographs (middle) of a bicistronic membrane-tethered CLIP^58^ (ORF1) probe containing a stop codon (red) and GFP (ORF2), expressed in HEK cells. Middle image show a higher power image of dashed region in left image. (**e**, right) Summary of the fraction of cells in which both ORFs could be detected in HEK (left) and HeLa (right) cells.

**Supplementary figure 4. Bicistronic expression of GFP and the potassium channel Kv4.2—a.** Illustration and distal sequence of Kv4.2 tagged with GFP (green, MVSK…), separated by UAA stop codon (red) and linker. **b.** Micrographs of cytoplasmatic expression of the post stop codon GFP in HEK cells. Higher power micrographh of the dashed area is shown on the right. **c.** Patch clamp recordings and IV curve of functional Kv4.2 necessarily located at membrane of cells.

**Supplementary figure 5. Effects of linker lengths and properties prior ORF2 on its expression in HEK cells— a.** Illustration of engineered clones; SNAP (gray) tethering BG-Alexa488 (green), stop codon (red), mCherry (magenta), partial sequence of the modified linkers (black), of the C terminus of the PDGFR (dark blue), stop codon (*, red), and the N terminus of mCherry (magenta). **b.** Micrographs of HEK cells surface labeled with BG-Alexa488 conjugated to membranal SNAP (green, left column), intracellular expression of mCherry (magenta, middle column). Linker types are noted next to micrographs: the original 12 aa long linker (‘original linker’), a short three aa linker (‘short linker’), a single nucleotide as linker causing a frame shift (‘frame shift’), without any linker (‘no linker’) and removal of the methionine from mCherry FP (‘no Met’). White arrows indicate cells with very low, albeit noticeable, expression of ORF2 (mCherry). **c.** Summary of fluorescence intensities of SNAP-tag (green) and mCherry (magenta channel, bottom), by cumulative distribution. Statistics are performed using a Kolmogorov-Smirnov comparison test. **d.** Frame shift enhances bicistronic expression. Fraction of cells expressing ORF1 only (green) and both ORFs (magenta).

**Supplementary figure 6. GEFRI clones enable polycistronic expression of three distinct ORFs separated by stop codons in HEK and HeLa cells— a.** Illustrations (left) of clone #1 (top), clone #2 (middle), and clone #3 (bottom) containing three ORFs: CFP (cyan), GCaMP7s (green) and tdTom (magenta). ORFs are solely separated by stop codons (red), w/o *Kozak* sequence. (right) Micrographs of variable expression of three ORFs in HEK cells. **b.** Normalized emission spectra of three colors obtained from expression of GEFRI_1_ (equally obtained for GEFRI_2_): CFP (dark blue), GCaMP7s (green) and tdTom (red). **c.** Percentage of cell showing expression of each ORF. The total sum exceeds 100%, as cells express multiple colors. Statistics were obtained by Two-way ANOVA, followed by *post hoc* Tukey multiple comparison test; **, p<0.01. **d.** Averaged responses of GCaMP7s from HeLa cells. Traces show average (bright green) and ± SEM (dark green) fluorescence responses (Δ*F/F*) following application of exogenous Ca^2+^ (50 µM), histamine (his. 10 µM, in case of HeLa cells), ionomycin (iono; 10 µM), and washing with EDTA for clone #1, clone #2 (n=30/each group).

**Supplementary figure 7. The importance of ORF1 for expression of following ORFs— a.** Illustration of the GEFRI_1_ clone in which we inverted the CFP (cyan, reversed CFP). **b.** Micrograph of HEK cells showing weak expression of all ORFs following transfection with the reversed clone (yellow arrowhead points to a cell that strongly expresses tdTom). **c.** Normalized emission spectra collected from cells from (**b**) in which we could detect fluorescence. Spectra was collected by excitation with 488 nm (blue line, green plot) and 561 nm (yellow line, magenta plot). Note the absence of any specific blue emission following excitation with 405 nm (blue area).

**Supplementary figure 8. Stable cells expressing GEFRI_2_ mainly exhibit two types of expression patterns— a.** Illustration of a HEK293t-stable line expressing GEFRI_2_ and micrographs (bottom) of cells showing expression of CFP (cyan; ORF1), GCaMP7s (green; ORF2) and tdTom (red; ORF3). **b.** Comparison of distribution of expression between different ORFs. 2D-plots show the various cellular populations (color-coded) based on expression levels of CFP and GCaMP (x axis-blue and green, respectively), then between GCAMP7s and tdTom (green and red), lastly between CFP and tdTom (blue and red). Fraction of each population is presented in (**c**) (colors correspond to those shown on FACS plots). **d.** Relationship between expression of ORFs. Spearman correlation was calculated for each two ORFs, shown as R^2^.

**Supplementary figure 9. Splitting GFP by a stop codon— a.** Sequences of GFP, sfGFP. The two parts of the original split-GFP (GFP_1-10_ and GFP_11_) and our split GFP (sGFP_1*M_) are annotated. **b.** Crystal structure of split-sfGFP (PDB: 2B3P, green). Splitting site is highlighted (yellow) and its sequence is noted. A stop codon (TGA, red) was introduced following the residue K214 (yellow), and the Met (gray) was frame-shifted by incorporation of a single bp. **c.** (top) Micrographs of HEK cells expressing GFP_1*M_ (left), GFP_1*Shift_, middle) or the first fragment of split GFP (isolated GFP_1-10_) (right). Arrowheads indicate specific green expressing cells; more easily discerned in bottom contrast-enhanced micrographs.

**Supplementary figure 10. The fragments of split-PA-sfGFP_2_ do not fluorescence on their own—**Micrographs of HEK cells showing complete lack of fluorescence when the fragments of split-PA-sfGFP_2_ are expressed alone, before or after photoactivation by 405 nm illumination. The first fragment includes the first 145 aa (PA-sfGFP_1-6_, top) and the second, 93 aa, (PA-sfGFP_7-11_, bottom). Micrograph of split-PAsfGFP_2_ prior photoactivation of dashed region (0 sec), post photoactivation (200 sec). Bottom right image shows 405 nm illumination of entire field of view (showing presence of monolayer of cells).

**Supplementary figure 11. Computational prediction of splitting sites for split GFP—** Snapshoot of the only splitting point predicted by SPELL (Split Protein rEassembly by Ligand or Light) computational approach^19^.

**Supplementary figure 12. Expression and functionality of Split-GCaMP6s variants— a.** Illustrations of split-GCaMP6s (based on PDB: 3WLC), and annotations of domains: M13 (purple), cpGFP (green), calmodulin (CaM; cyan), stop codons (red filled stars-sites of insertion of a stop codon, whereas empty red star indicate replacement of an extant residue in GCaMP6s by a stop codon). Sequences of interest are noted above site within the probe. **b.** Micrograph of HEK cells expressing the clones presented in (**a**), prior (left) and following (right) application of 10 µM ionomycin (-/+ Iono). Summary of maximal responses (Δ*F/F*) are presented on the right. **c, d.** Micrographs of hippocampal neurons following transfection with GCaMP6s_3*_ (**c**) or M13-cpGFP (**d**) alone (top micrographs) or co-expressed with another plasmid encoding CaM (+CaM). Traces show representative responses for each instance (traces are color coded as arrowheads in micrographs), **e.** Summary of maximal Δ*F/F* for GCaMP6s_3*_ and M13-cpGFP, with (black) or without (grey) co-expression of CaM in cultured hippocampal neurons. **f.** Micrograph of simultaneous electrical (loose patch, black trace) and fluorescent recordings (color traces) of a pyramidal neuron co-expressing M13-cpGFP and CaM. Ca^+2^-responses were recorded by a standard camera and LED illumination source of an electrophysiology set-up. Traces represent activity recorded from various locations of the cell, as indicated by colored arrowheads in micrograph. The numbers below black trace indicate the number of action potentials for each event. Note that the probe can report on single action potentials. **g.** Decay time (left) and the Δ*F/F* (right) as a function of number of action potentials. **h, i.** Micrographs of representative hippocampal neurons following single or double infections with AAV M13-cpGFP (**h**), or co-infection with AAV-CaM (**g**). Traces shown a variety of ca^2+^-responses. Summary is shown in (**j**). Statistical significance was tested by multiple comparisons of mixed-effects analysis (**b**), and non-parametric Mann-Whitney test (**e** and **j**).

**Supplementary figure 13. Optimization of split GCaMP_2_— a.** Illustrations of SPLIT superfolder (sf)- and optimal superforlder (osf)- GCaMP6s_2_ variants. Optimizations mutations are denoted (grey stars). **b.** Micrographs of HEK cells expressing the clones presented in (**a**), prior (left panels) and following application of 10 µM ionomycin (+Iono) (right). **(c)** Summary of Ca^+2^- depented fluorescence change (Δ*F/F*). Dashed violin plots are taken from **Supplementary figure 12b** to ease comparison between variants. **d.** Individual fragemtns from split-GCaMP6s2 are not fluorescent. Micrographs of HEK cells expressing the fragments of split-GCaMP6s_2_ individually (as noted atop micrographs) show no fluorescence. Significances were tested by multiple comparisons of mixed-effects analysis.

**Supplementary figure 14. Transferability of splitting site to a red-GECI— a.** Illustrations of split jR-GECO1a, including its fragments; M13 (purple), cp-mApple (red), calmodulin; CaM (cyan), and splitting localization sites. **(b)** Micrographs of HEK cells expressing the jR-GECO1a; prior and following application of 10 µM ionomycin (+Iono); summarized on the right. Significance was tested by multiple comparisons of mixed-effects analysis.

**Supplementary figure 15. Using novel split-GFP to highlight mitochondria-ER contact sites—a.** Illustration of the split GFP variants employed to label mitochondria (TOM20-GFP1-6) and ER. GFP_1-6_ (green) tethered to TOM20 (yellow) and GFP_7-11_ (green) tethered to ER membrane by CB5 (orange) and mRuby_3_ (red), to express at mitochondria and ER, respectively (as depicted in cartoon). **b.** Micrograph of TOM20-GFP_1-6_ (part I; top) and GFP_7-11_-CB5-mRuby_3_ (part II; bottom) expression in HEK. Note the complete absence of green fluorescence when the fragments are not expressed jointly, while ER can be clearly discerned by mRuby_3_ (magenta) **c.** Micrographs of HEK cells co-expressing TOM20-GFP_1-6_ (top left panel; green) and the GFP_7-11_-CB5-mRuby_3_ (top middle panel; magenta). (bottom) A high power image (of dashed region in top right micrograph) showing overlap between green and magenta, summarized in fluorescence profile plots (bottom right). Note the imperfect overlap, as magenta does not depend on the expression of both fragments, whereas green does. Green fluorescence does not appear to highlight the structure of mitochondria, rather provides localized puncta with ER. **d.** TOM20-GFP_1-6_ is expressed across the entire mitochondrial-membrane. (left) Projection micrographs of HEK cells co-expressing the mitochondrial TOM20-GFP_1-6_ probe and a soluble GFP_7-11_ fragment, as well as mitochondrial ca^2+^ indicator, mito-CAR-GECO1 (magenta). Note the high number of labeled mitochondria and overlap with magenta. (Inset) high power images for the dashed yellow region, showing green fluorescence (at membrane of mitochondria) surrounding magenta (expressed within mitochondria). These are shown in small fluorescence profile plots.

**Supplementary figure 16. MEGIC_1_—engineering a <u>M</u>ito-<u>E</u>R <u>G</u>enetically encoded <u>I</u>ndicator for <u>C</u>alcium— a.** MEGIC_1_ is composed of TOM_20_-M13-cpGFP and CaM-CB5-mRuby_3_ (mitochondrial and ER expression, respectively). Micrograph of a HeLa cell co-expressing TOM_20_-M13-cpGFP and mito-CAR-GECO1. Note the almost perfect overlap, as noted in **Suppl. 15d**. **c.** MEGIC1 reports on Calcium activity as the mitochondrial mito-CAR-GECO1 probe. HeLa co-expressing TOM_20_-M13-cpGFP and mito-CAR-GECO1, and traces from individual cells on the right. (magenta-pink traces represent activity of mito-CAR-GEWCO1, whereas light and dark green traces represent concomitant activity monitored by MEGIC1). **c, d.** Micrographs of HEK cells expressing various combination of MEGIC1 clones (**a**), prior (left panels) and following application of 10 µM ionomycin (+Iono); summarized in (**d**).

**Supplementary figure 17. MEGIC_1_ in neurons *in vitro* and *in vivo—* a, b,** Micrographs of infected pyramidal neurons expressing only the first fragment (TOM20-M13-cpGFP, **a**) or both fragments (**b**) of MEGIC_1_, and corresponding Ca^2+^-responses (Δ*F/F*) shown on the right (colored traces corresponding to arrowheads in micrographs; summarized in (**c**). **d, e,** Two representative examples of neurons co-expressing both fragments of MEGIC_1_, dually patched (patch pipette, dashed white indication in micrographs) and imaged; corresponding traces on the right (color-fluorescence, black-electrical activity, whole cell mode). Insets-zoom-in onto electrical and optical activities from region in traces (dashed black regions). **f.** Summary of maximal Δ*F/F* obtained in soma (top) and processes (below), and correspondence towards #APs (n = 6). Significances were tested by multiple comparisons of mixed-effects analysis. g. Micrographs of a Zebrafish larva expressing both fragments, with high power micrographs shown on the right, and activity for select regions (color coded as arrowhead sin micrograph); summarized in violin plot (bottom right).

**Supplementary figure 18. Engineering an improved MEGIC reporter *for* neurons— a.** Depiction of the new splitting site in MEGIC_2_. **b.** Micrographs of HEK cells expressing individual fragments of standard-, superfolder (sf)- or optimal superforlder (osf)-MEGIC_2_. Note lack of fluorescence in green channel, whereas magenta (demarking ER) is noticeable when second part is expressed, summarized in (**c**). **d.** Micrographs of cultured neurons infected with individual viruses encoding each of the fragments of sf-MEGIC_2_. **e.** Monitoring electrical and fluorescent activity jointly. Traces show robust intracellular ca^2+^-activities (colored traces) without any AP firing (black bottom trace) by neuron.

**Supplementary figure 19. ER stress affects expression stoichiometry of ORFs in GEFRI_2_—a.** Established mechanism of tunicamycin (TM). **b.** ER stress enhances the relationship (joint expression) between the various ORFs in a dose-dependent manner. **c.** TM engenders strong upregulation in expression of second and third ORFs, as indicated by the changes in the populations, highlighted by colored bars below. **d.** Summary of expression levels. Plot represent all pixel cumulative distribution (normalized frequency vs. pixel intensity). Correlations between two independent parameters (R^2^) were examined using Spearman correlation, and cu ulative distributions were examined by Kolmogorov-Smirnov comparison tests. Asterisks indicate statistically significant differences as follows: *, P < 0.05; **, P < 0.01; ***, P < 0.001.

## Notes

### Competing Interest Statement

The authors have declared no competing interest.

