## Supplementary figures and images for "A novel polycistronic method tailored for engineering split GECIs"

### supplementary figure 1

#### Olszakier and Hussein et al. Supplementary Figure 1

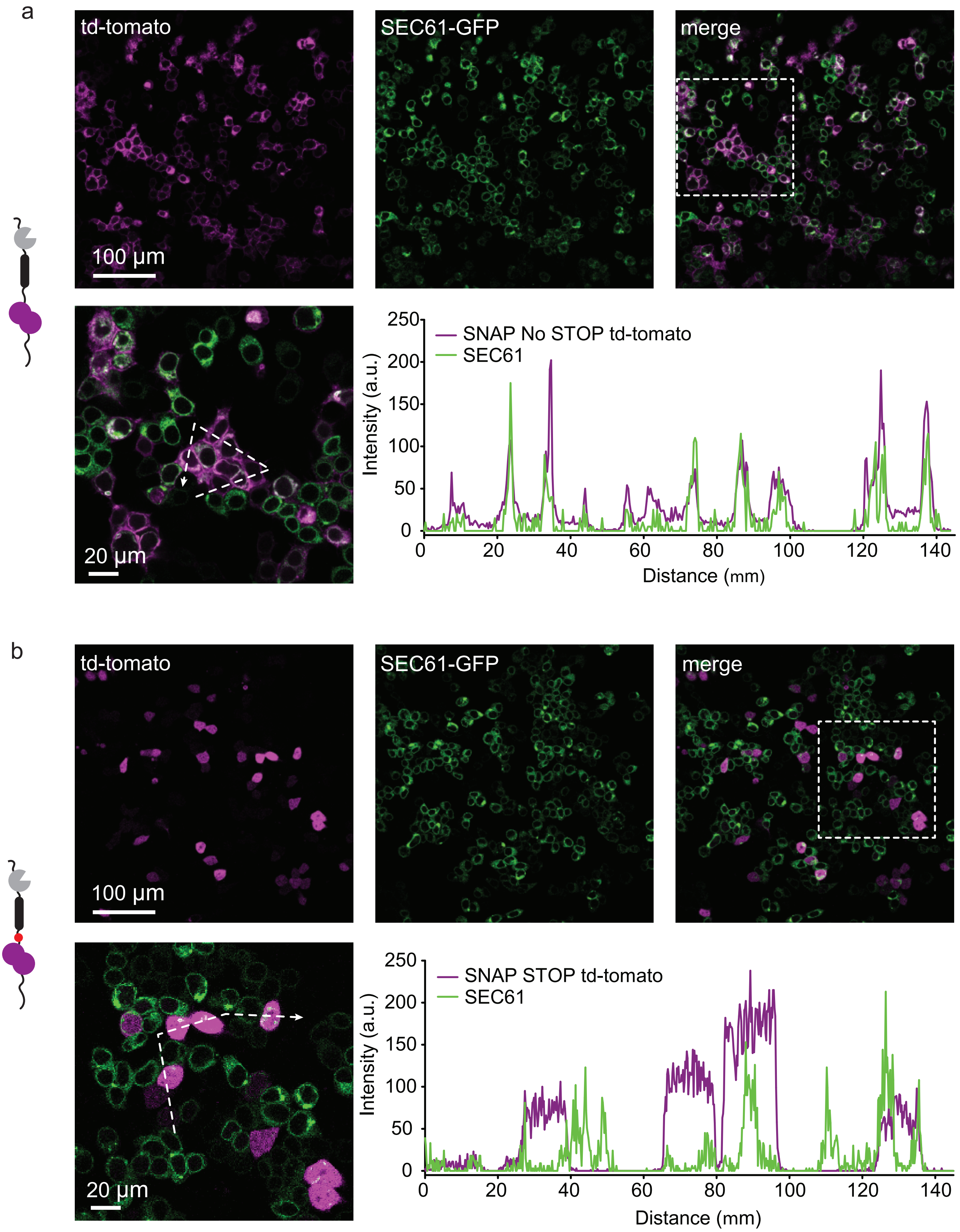

### supplementary figure 2

Olszakier and Hussein et al. Supplementary Figure 2

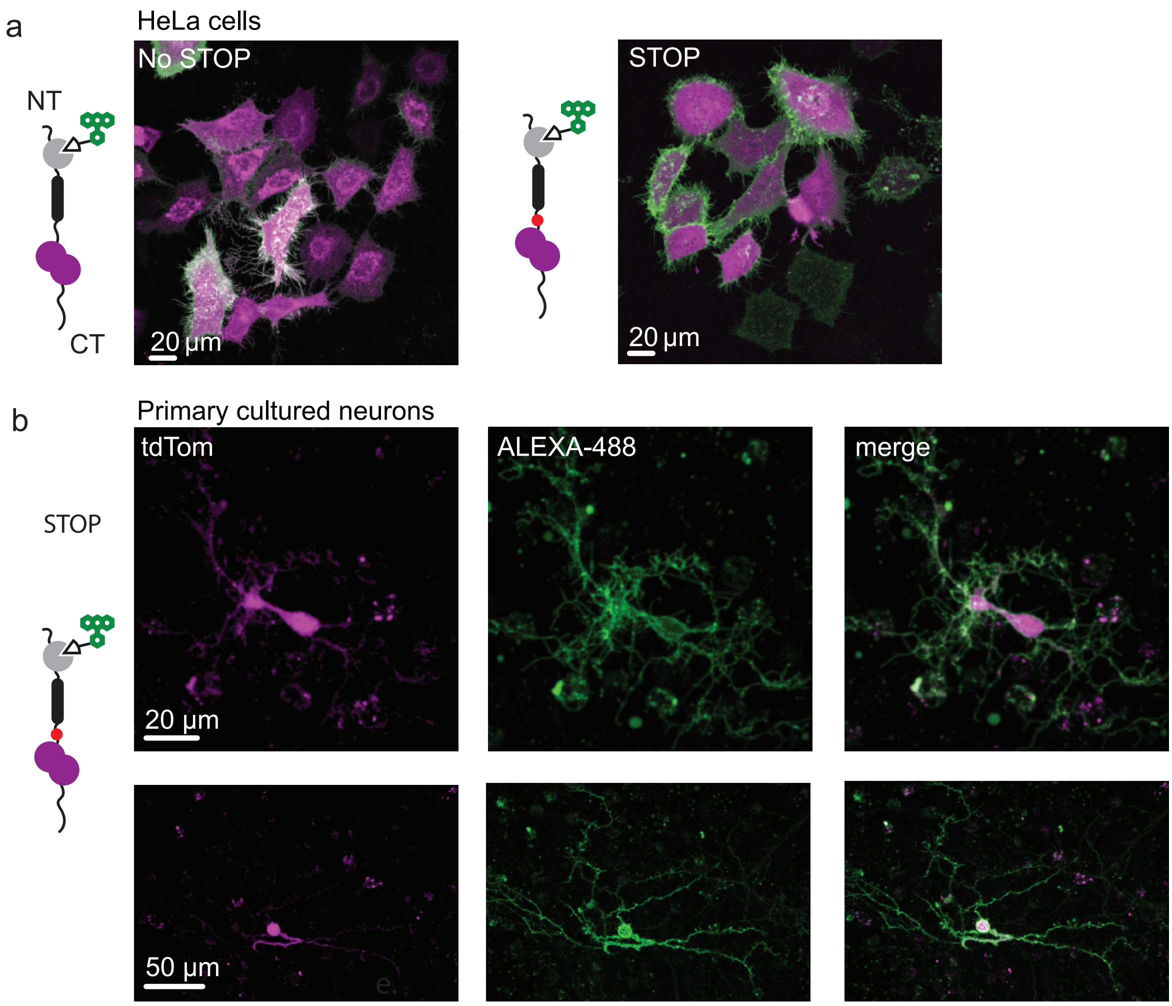

### supplementary figure 3

Olszakier and Hussein et al. Supplementary Figure 3

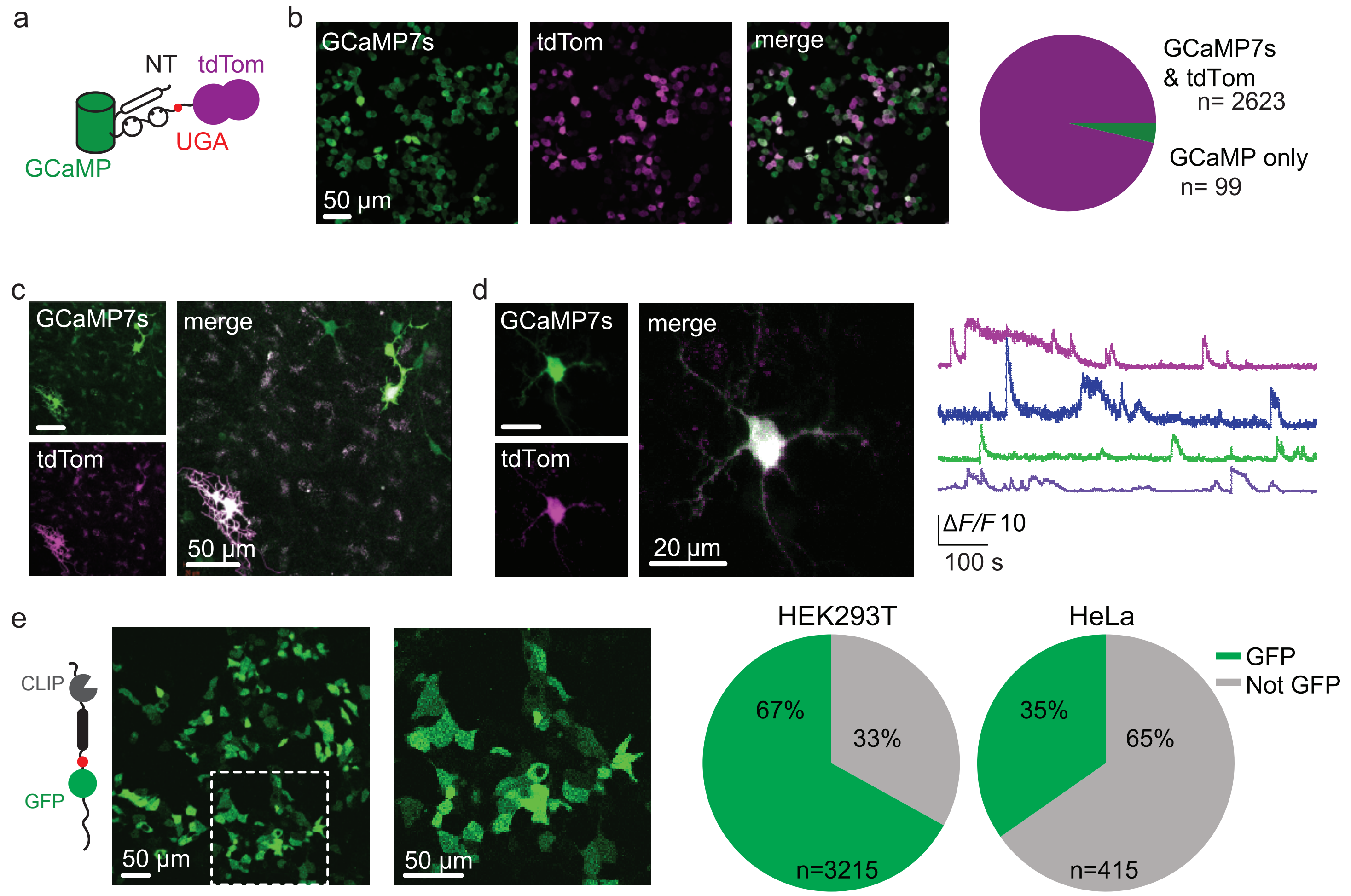

### supplementary figure 4

Olszakier and Hussein. Supplementary Figure 4

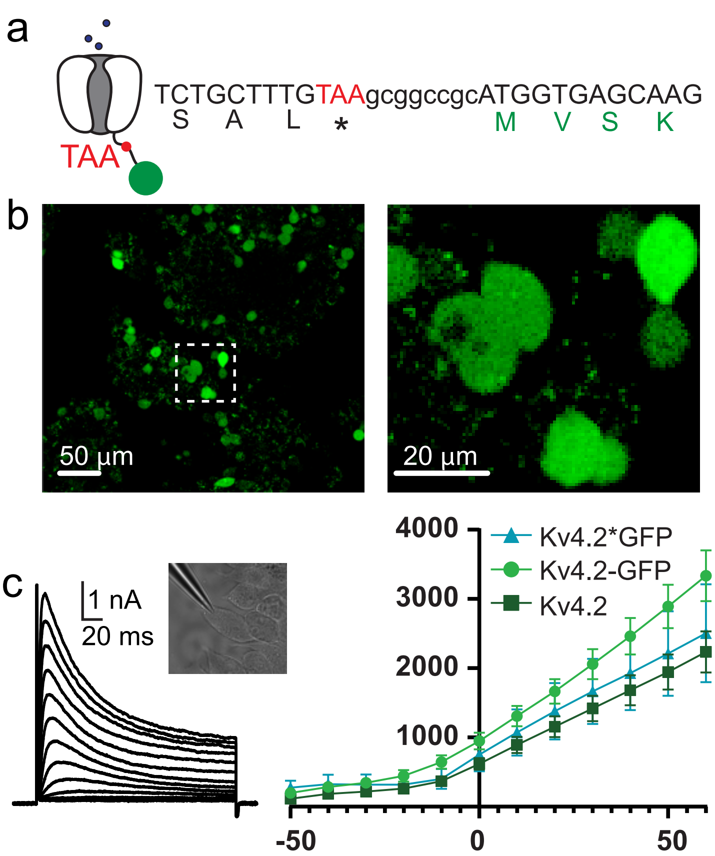

### supplementary figure 5

#### Olszakier and Hussien et al. Supplementary Figure 5

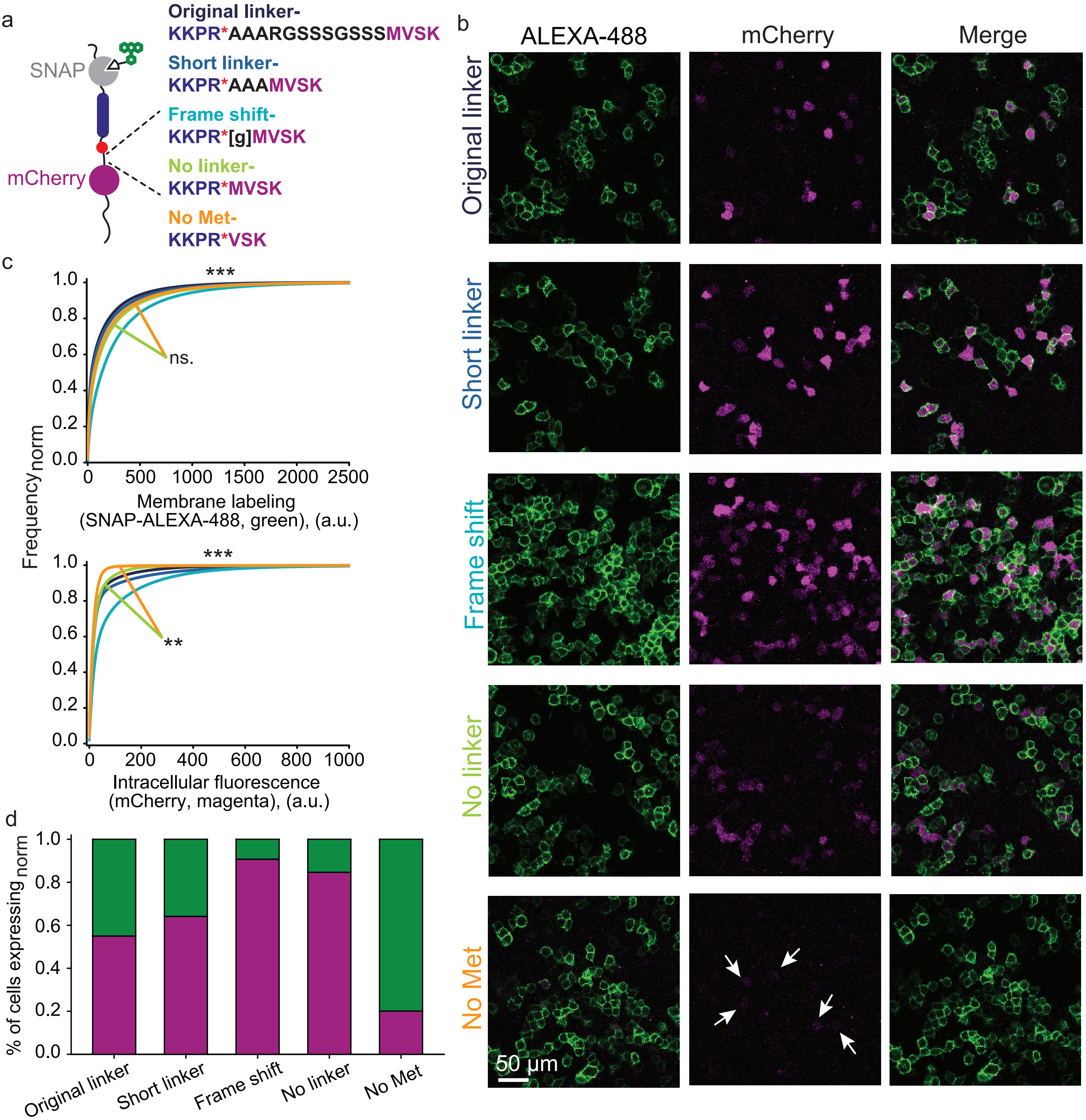

### supplementary figure 6

#### Olszakier and Hussein et al. Supplementary Figure 6

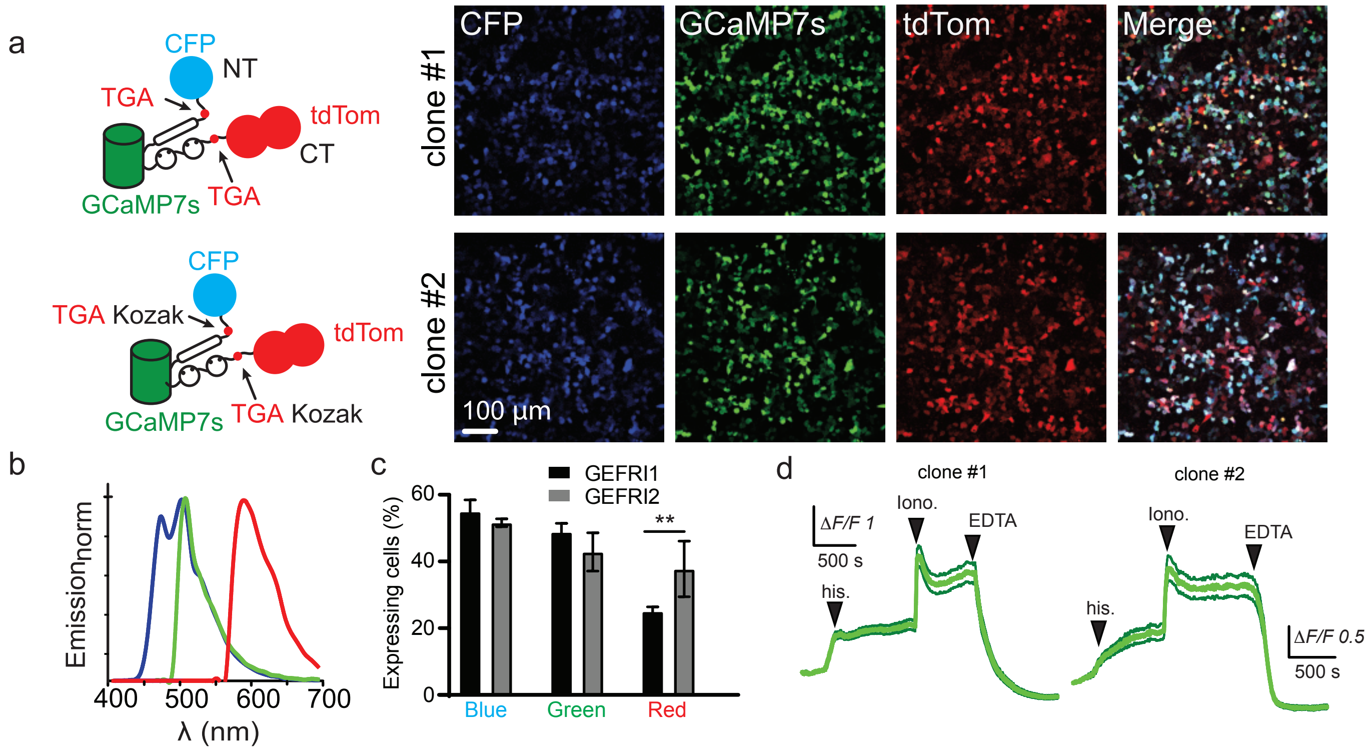

### supplementary figure 7

#### Olszakier and Hussein et al. Supplementary Figure 7

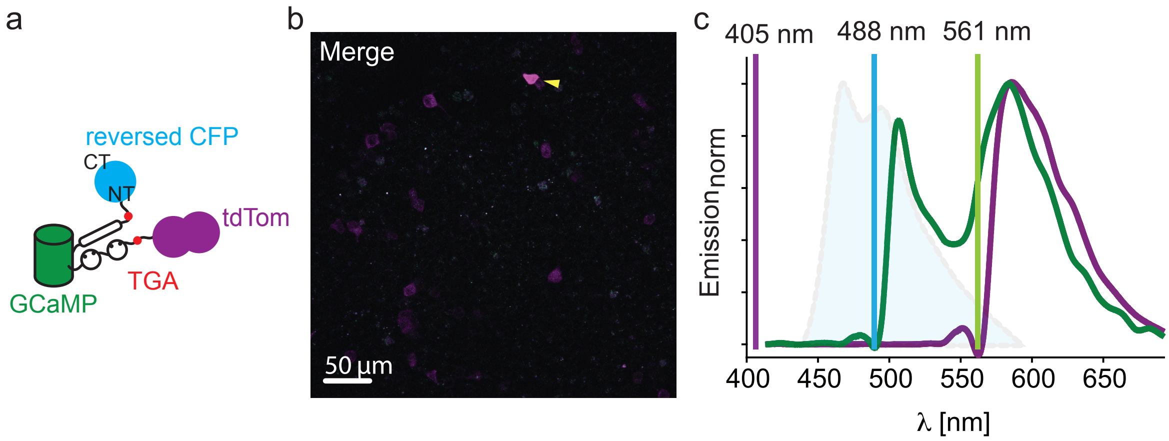

### supplementary figure 8

Olszakier and Hussein et al. Supplementary Figure 8

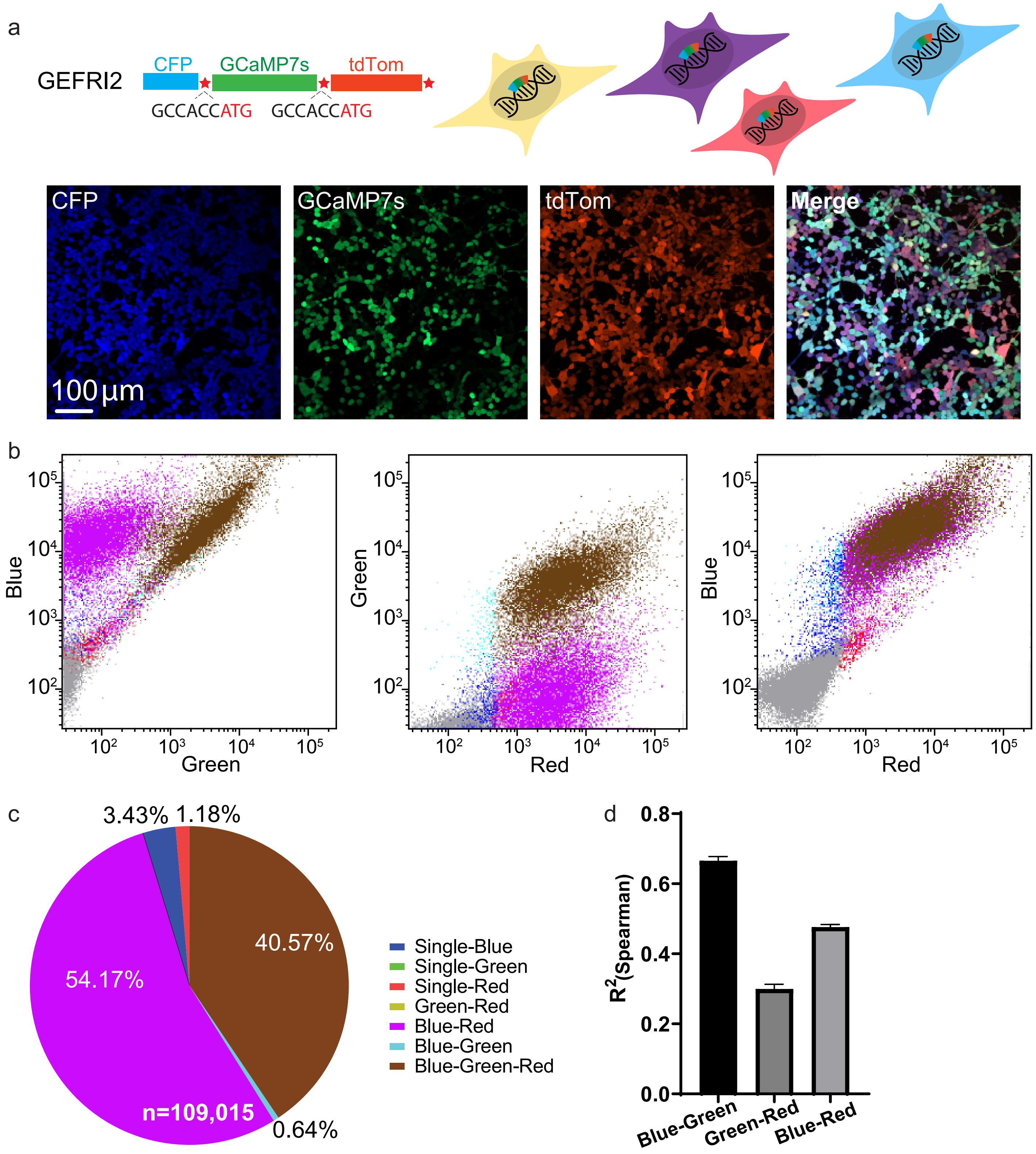

### supplementary figure 9

#### Olszakier and Hussien et al. Supplementary Figure 9

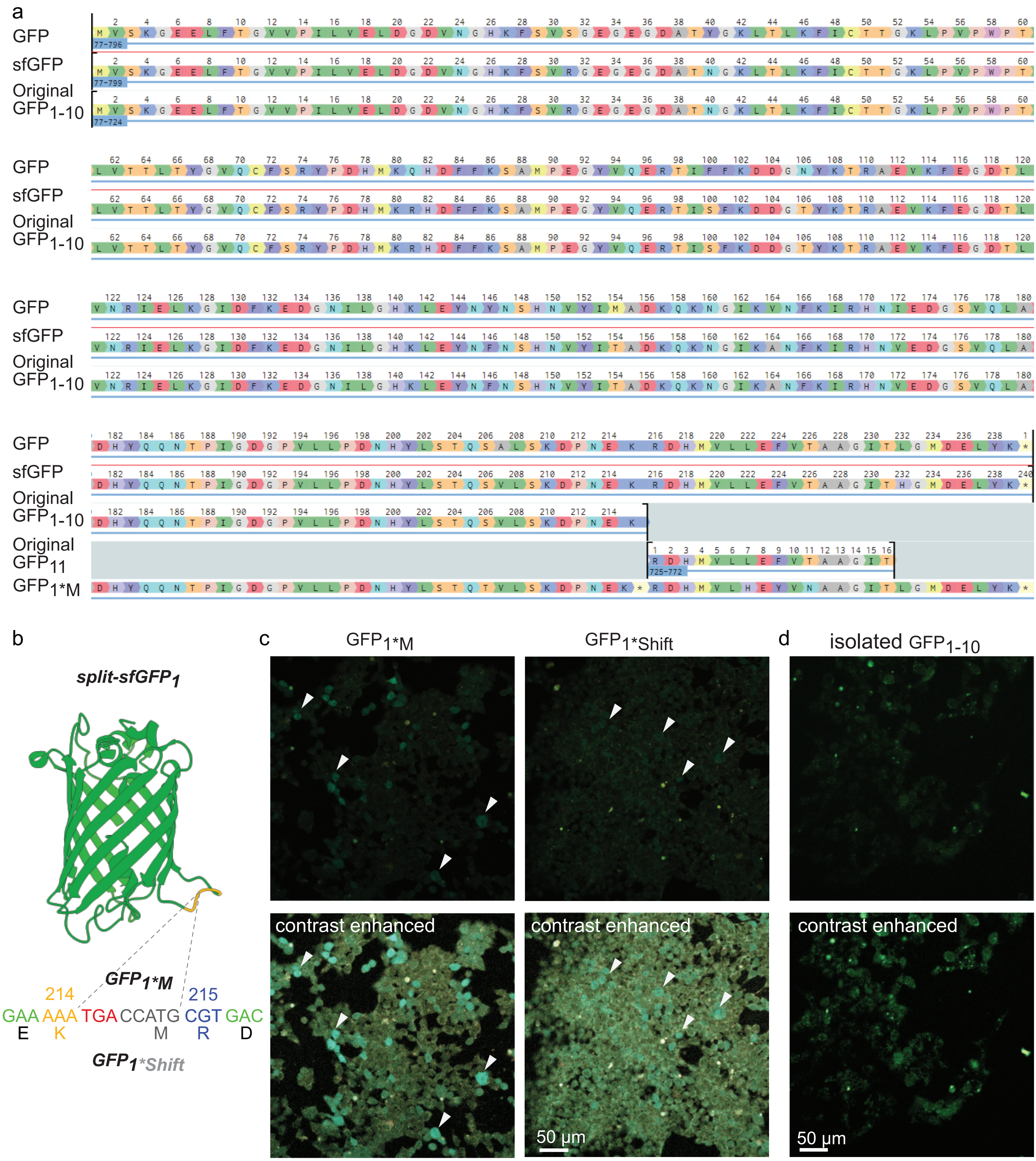

### supplementary figure 10

#### Olszakier and Hussien et al. Supplementary Figure 10

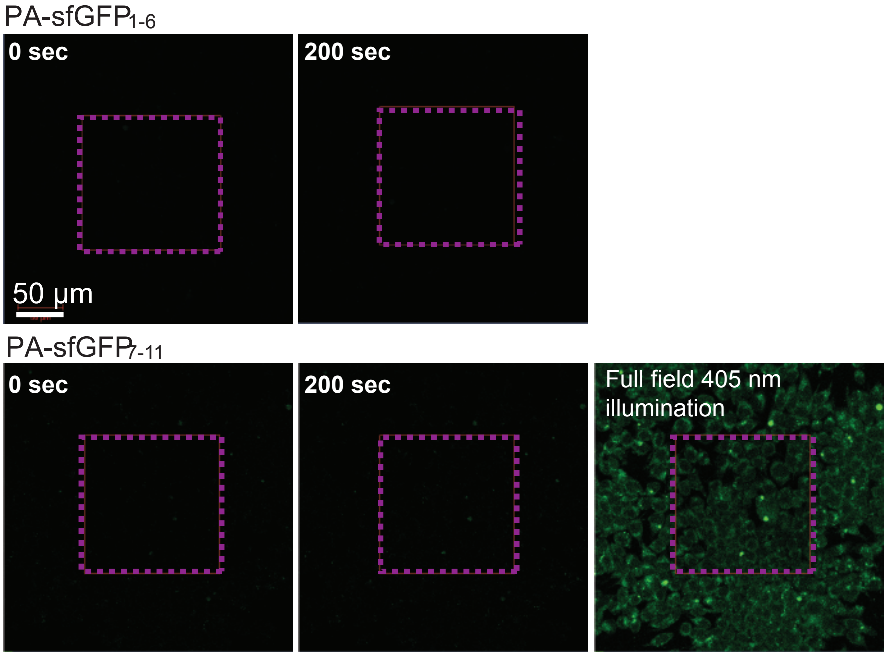

### supplementary figure 11

Olszakier and Hussien et al. Supplementary Figure 11

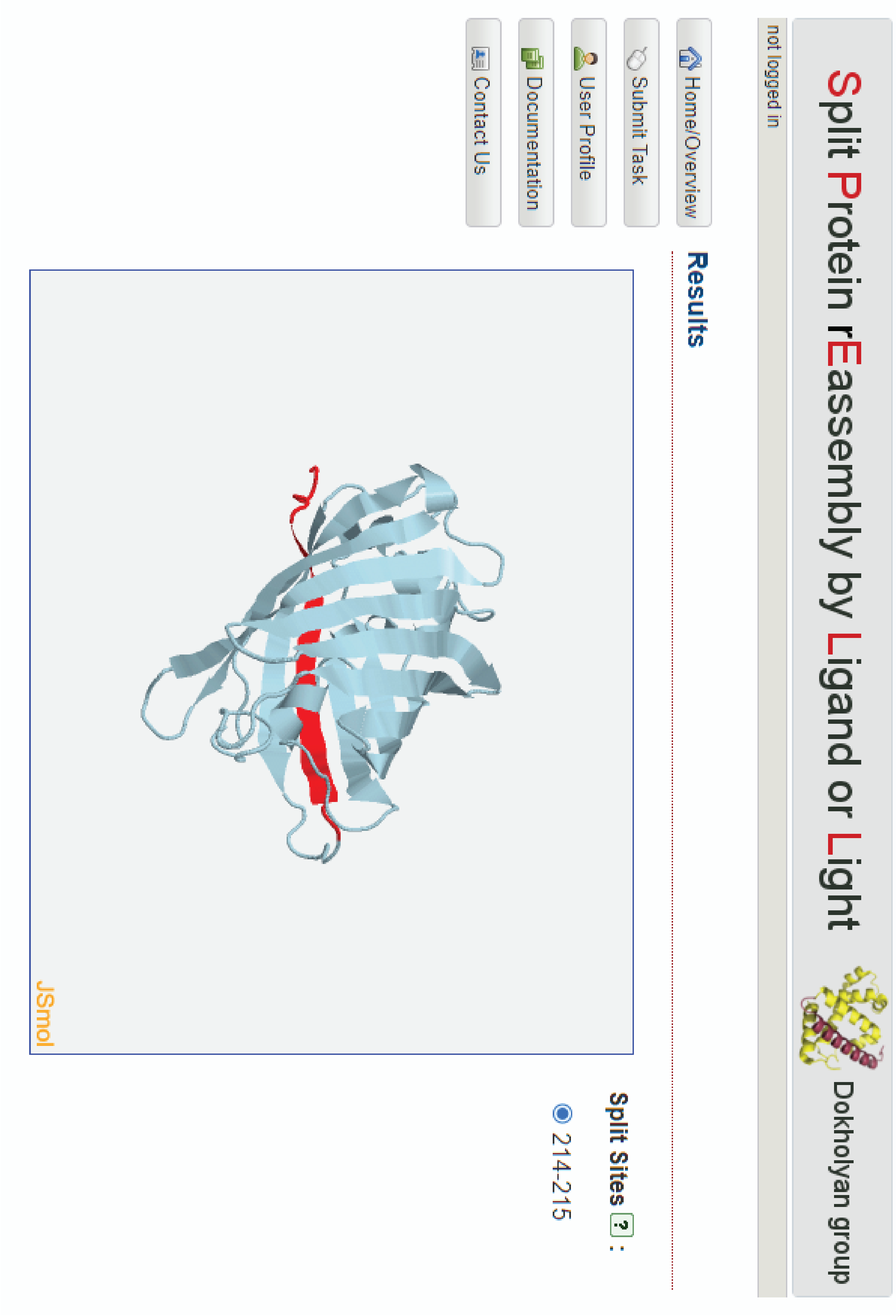

### supplementary figure 12

#### Olszakier and Hussien et al. Supplementary Figure 12

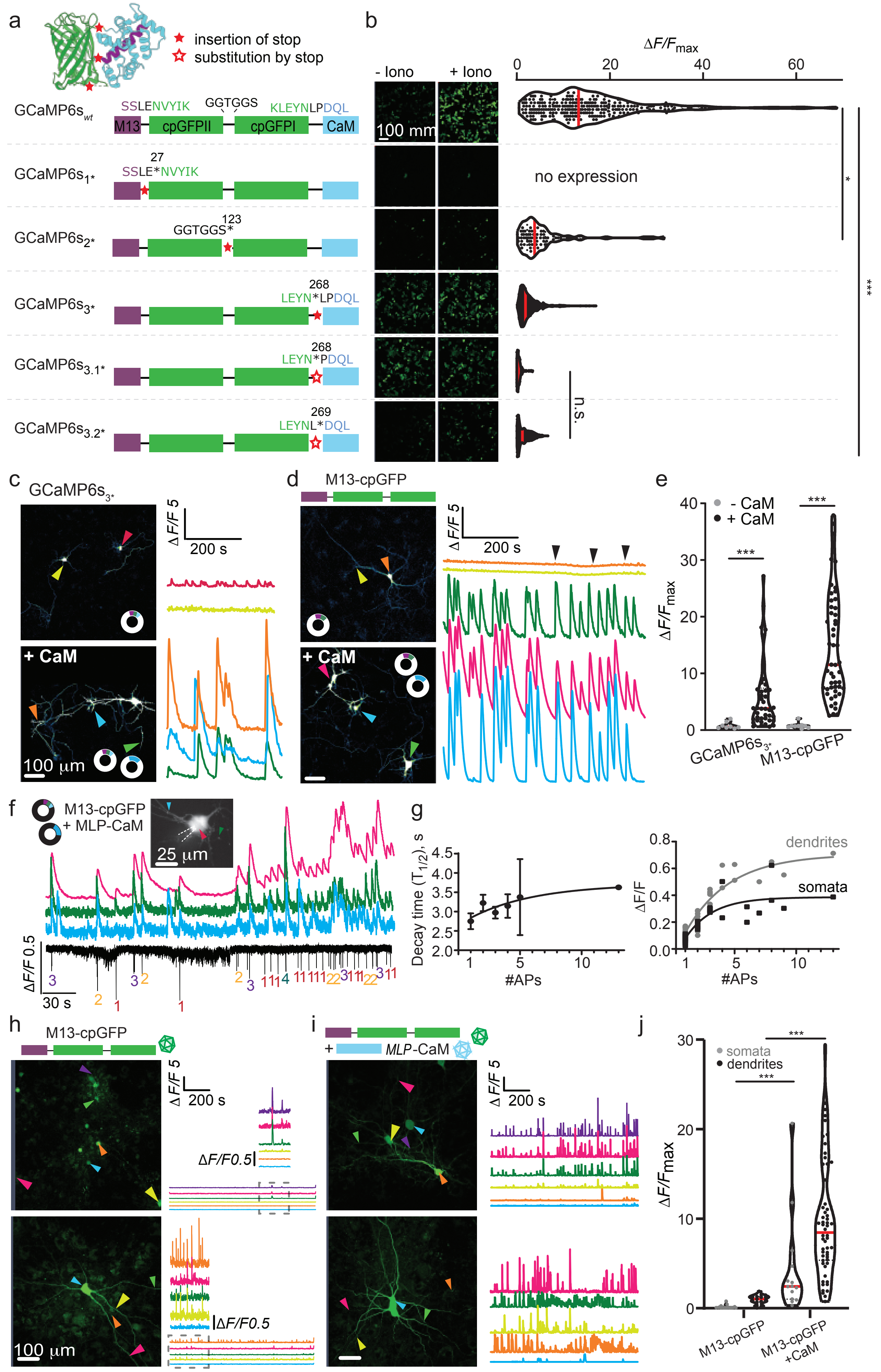

### supplementary figure 13

Olszakier and Hussien et al. Supplementary Figure 13

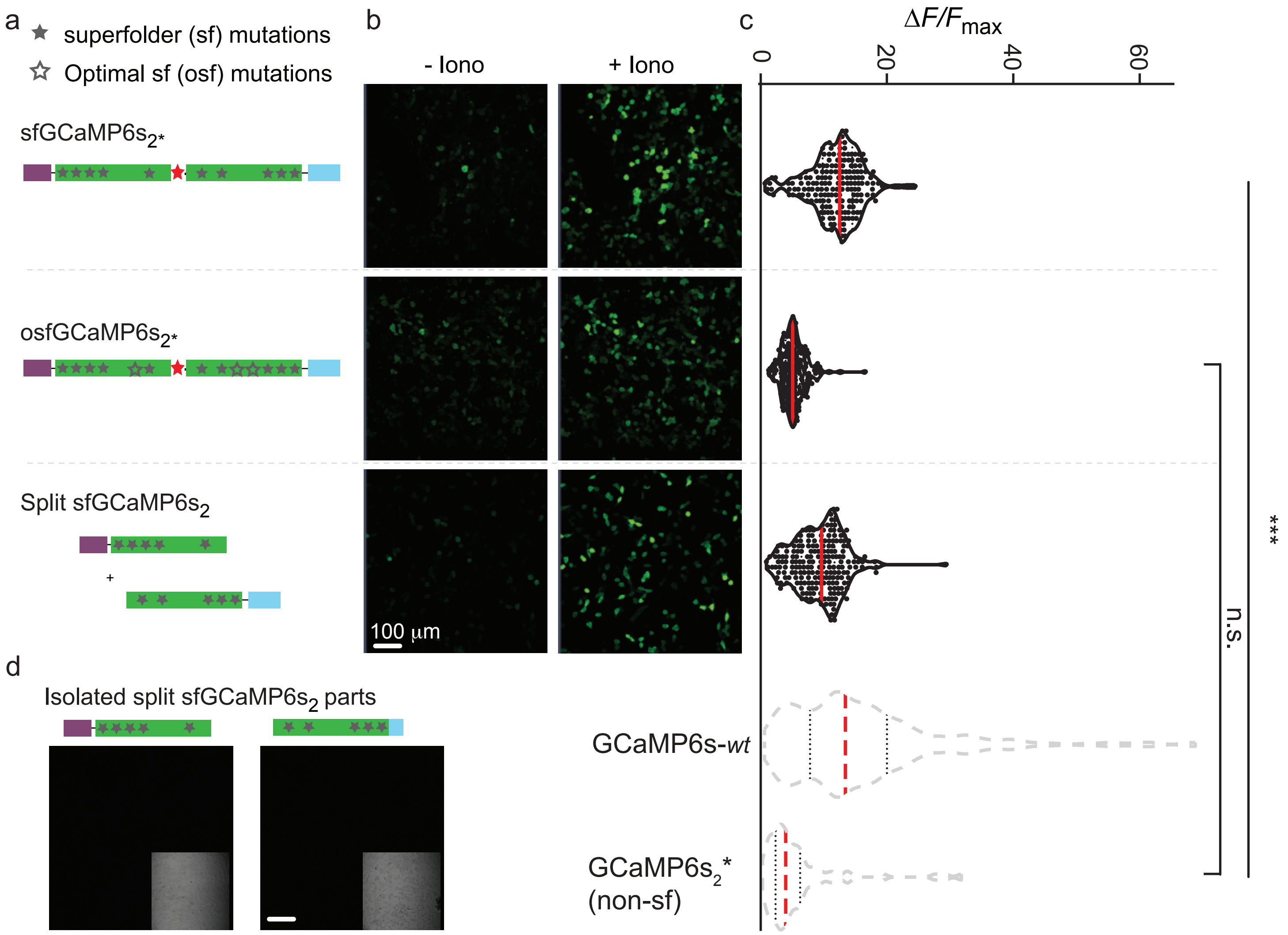

### supplementary figure 14

Olszakier and Hussien et al. Supplementary Figure 14

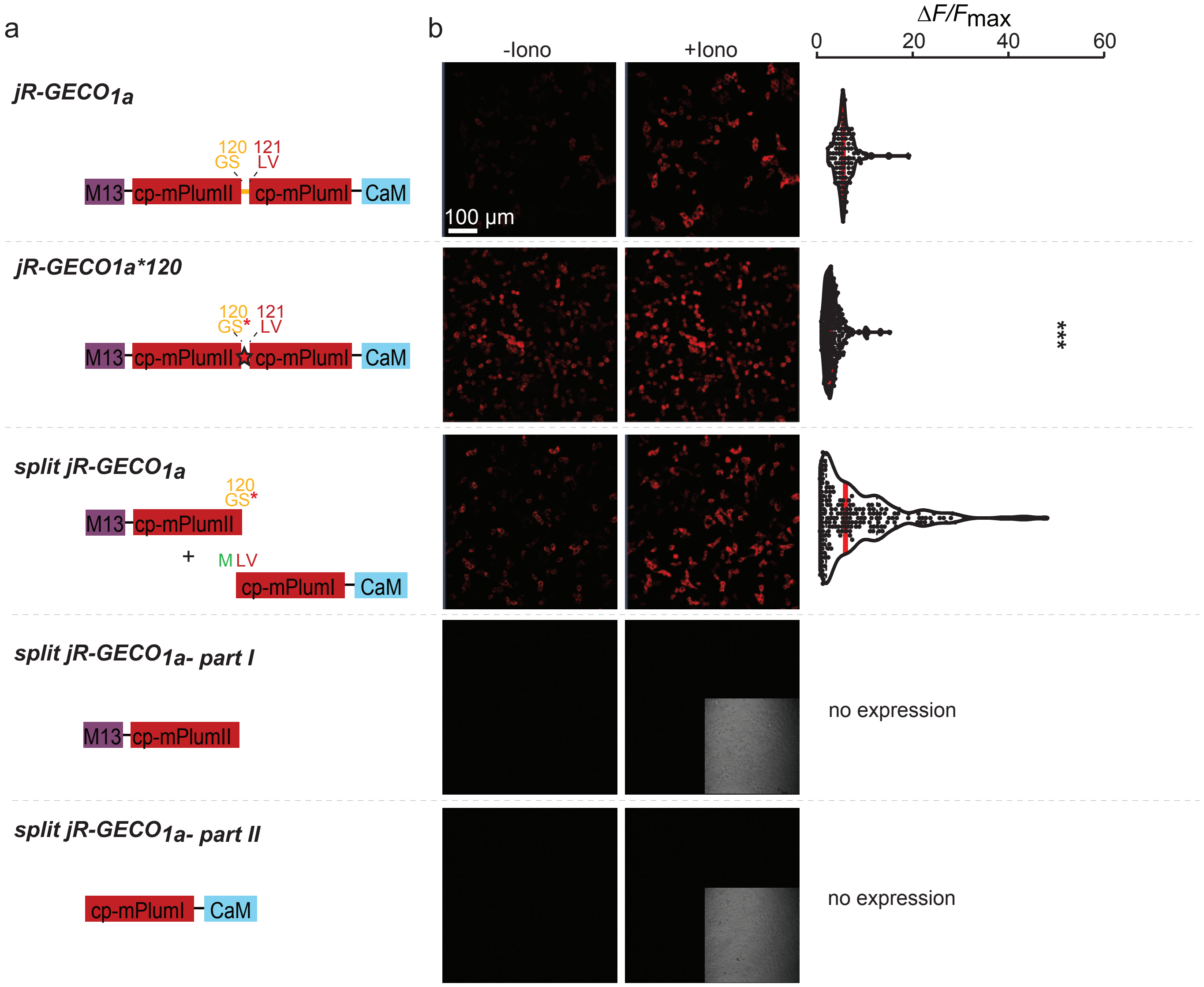

### supplementary figure 15

Olszakier and Hussien et al. Supplementary Figure 15

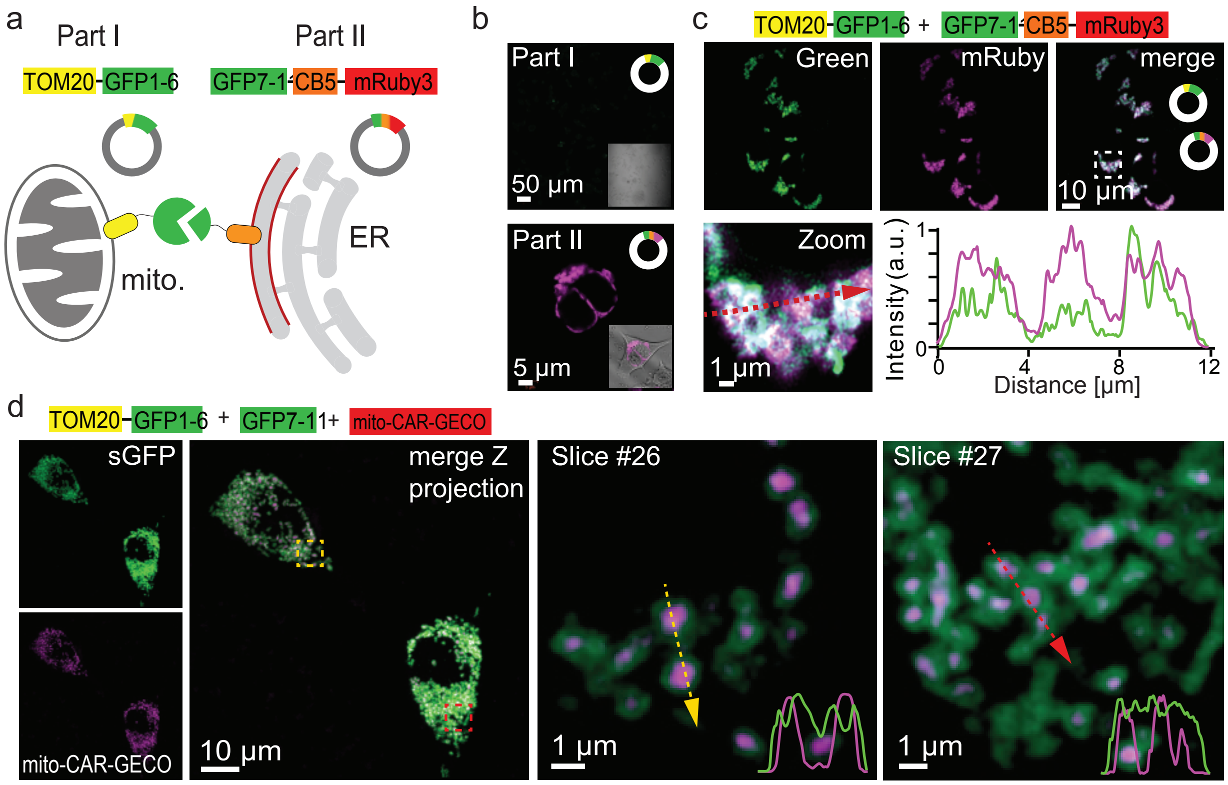

### supplementary figure 16

Olszakier and Hussien et al. Supplementary Figure 16

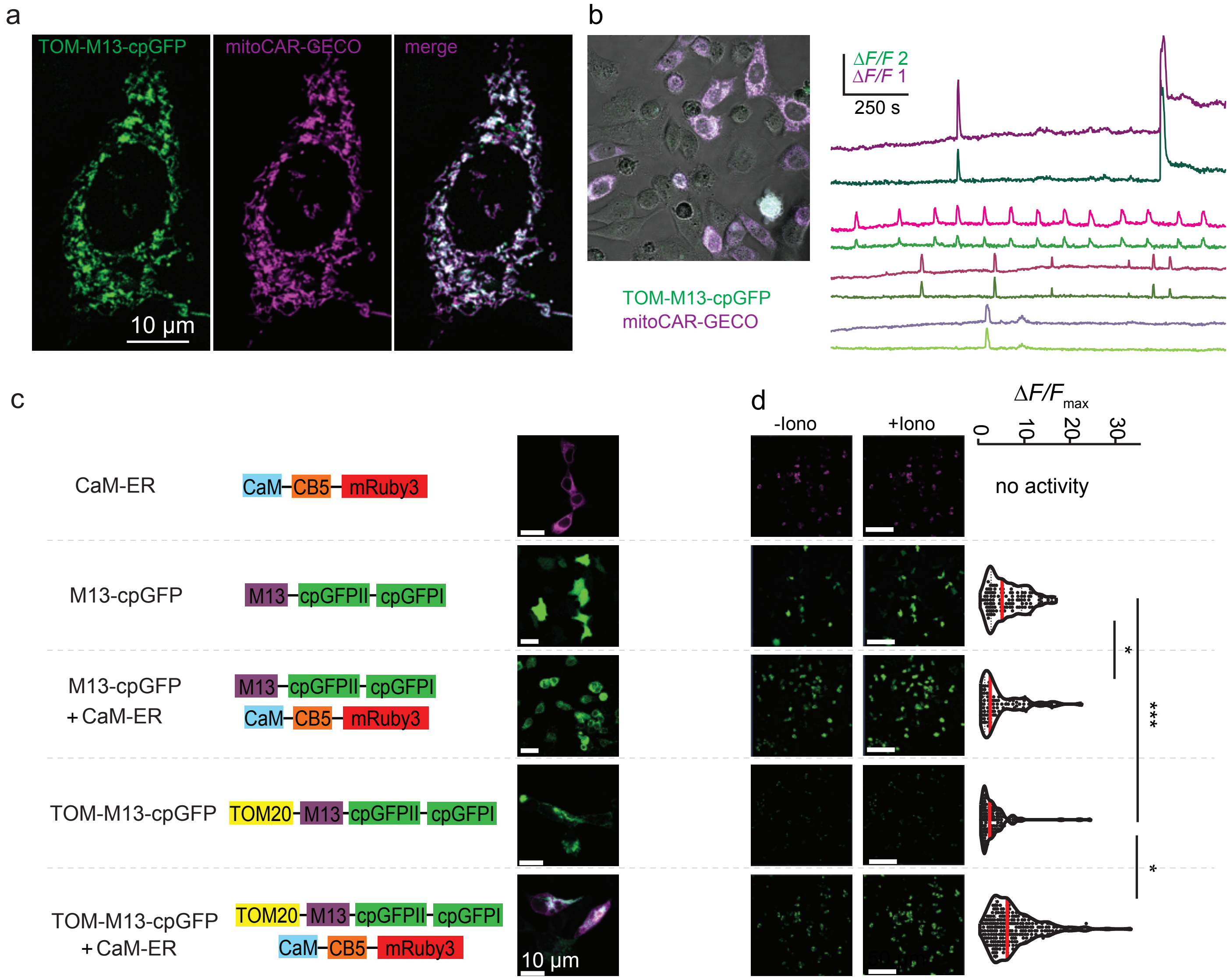

### supplementary figure 17

Olszakier and Hussien et al. Supplementary Figure 17

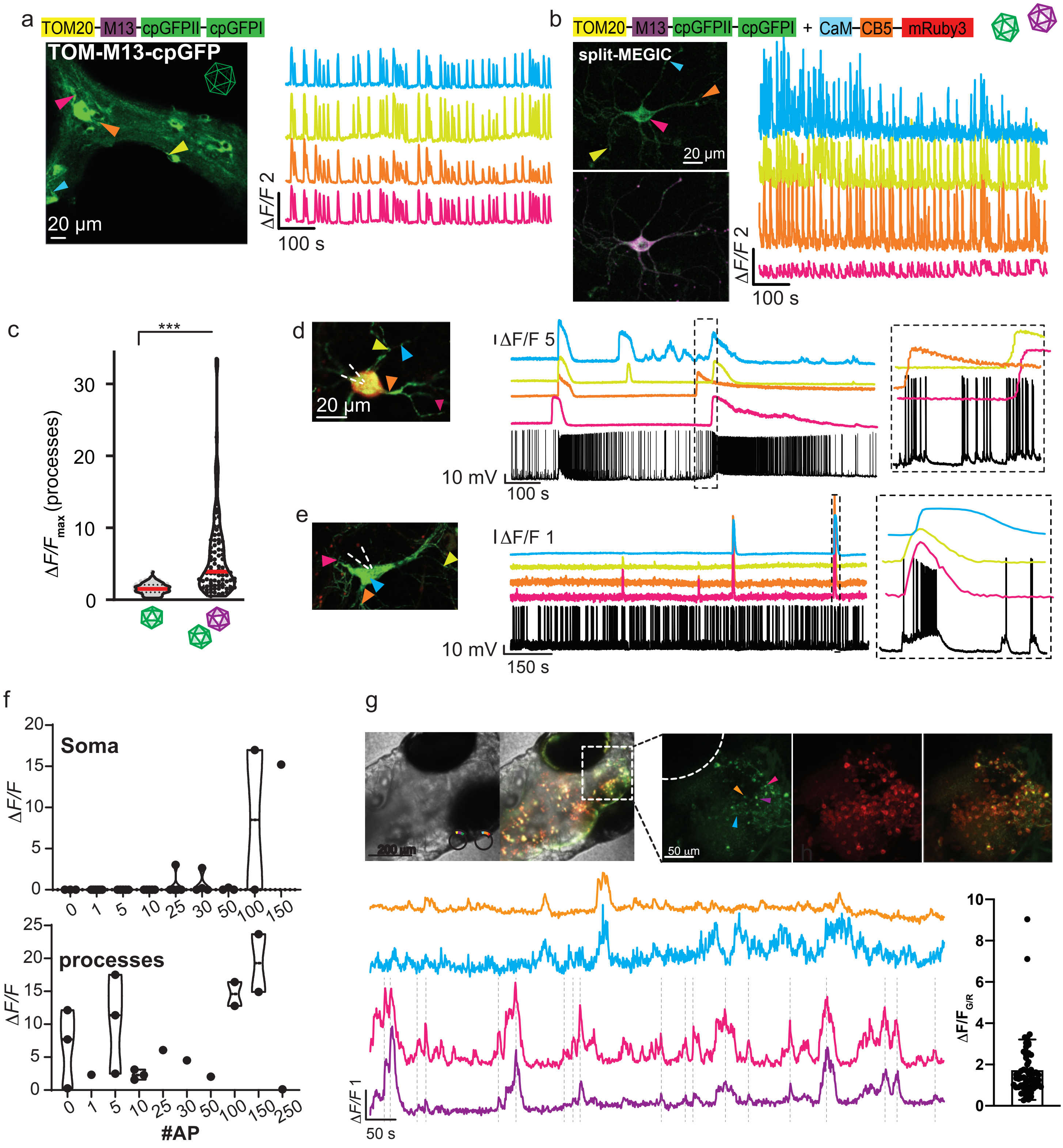

### supplementary figure 18

Olszakier and Hussien et al. Supplementary Figure 18

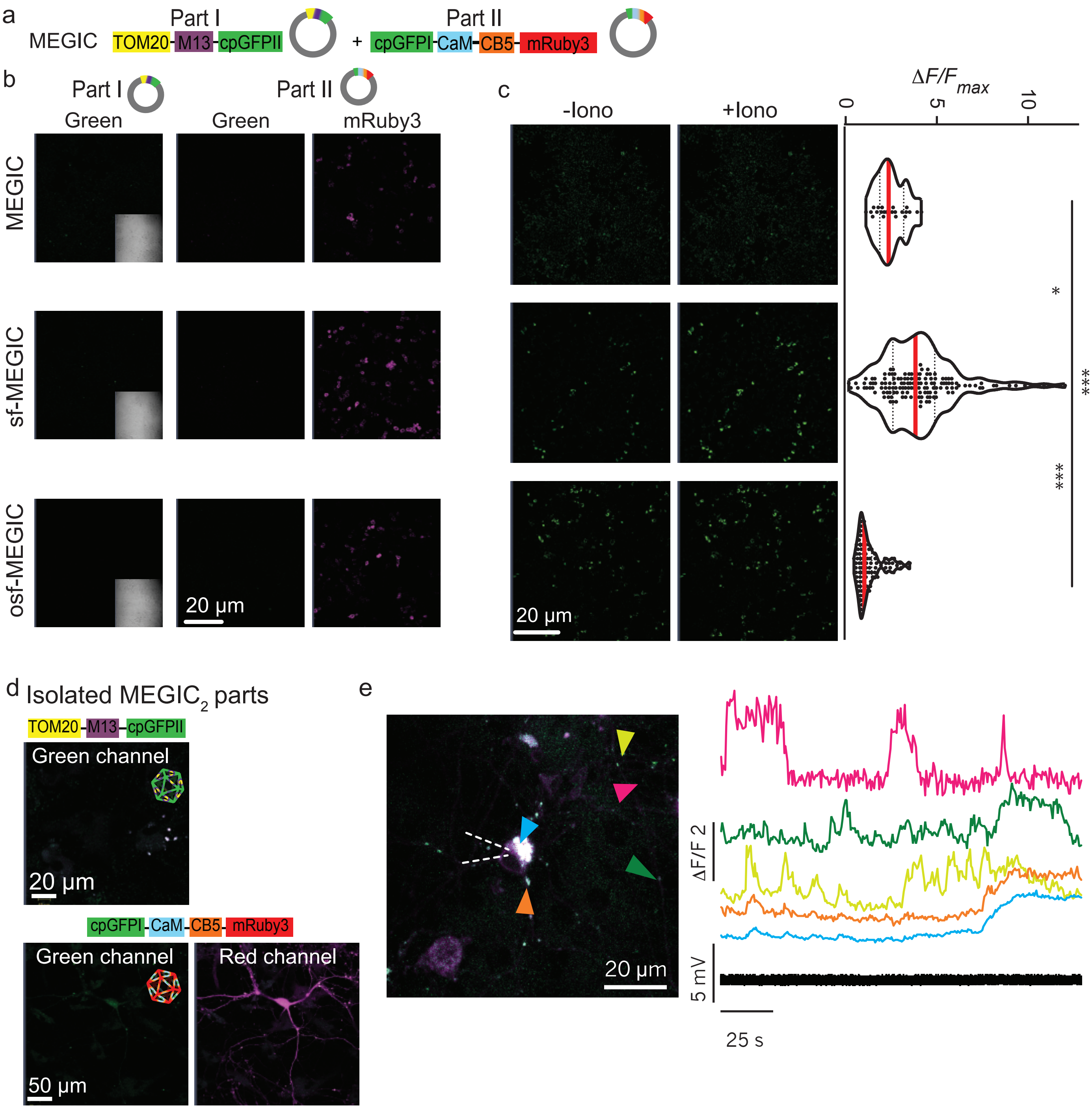

### supplementary figure 19

#### Olszakier and Hussien et al. Supplementary Figure 19

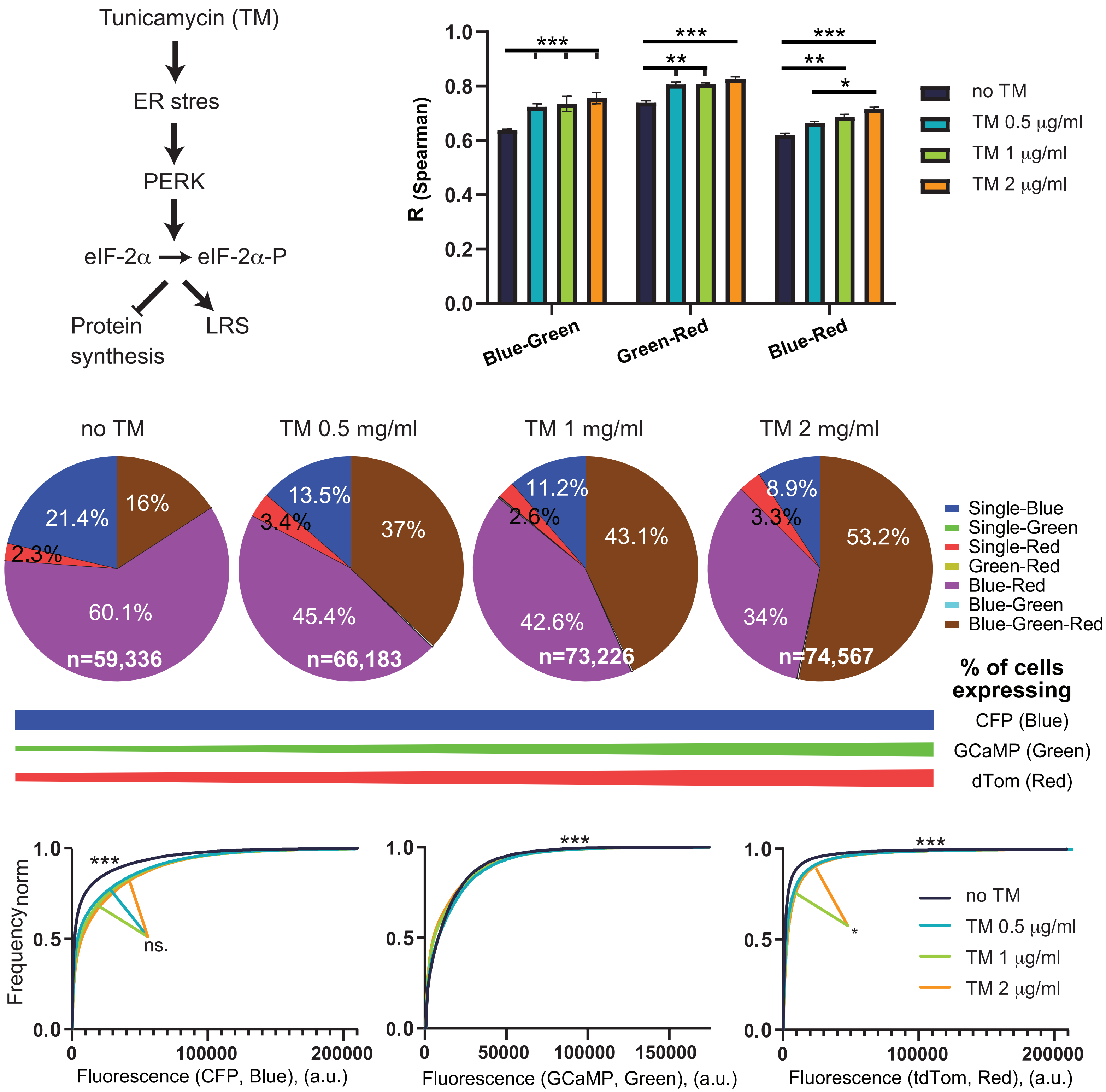
